# An antibody targeting the N-terminal domain of SARS-CoV-2 disrupts the spike trimer

**DOI:** 10.1101/2022.01.12.476120

**Authors:** Naveenchandra Suryadevara, Andrea R. Shiakolas, Laura A. VanBlargan, Elad Binshtein, Rita E. Chen, James Brett Case, Kevin J. Kramer, Erica Armstrong, Luke Myers, Andrew Trivette, Christopher Gainza, Rachel S. Nargi, Christopher N. Selverian, Edgar Davidson, Benjamin J. Doranz, Summer M. Diaz, Laura S. Handal, Robert H. Carnahan, Michael S. Diamond, Ivelin S. Georgiev, James E. Crowe

## Abstract

The protective human antibody response to the severe acute respiratory syndrome coronavirus 2 (SARS-CoV-2) virus focuses on the spike (S) protein which decorates the virion surface and mediates cell binding and entry. Most SARS-CoV-2 protective antibodies target the receptor-binding domain or a single dominant epitope (‘supersite’) on the N terminal domain (NTD). Here, using the single B cell technology LIBRA-seq, we isolated a large panel of NTD-reactive and SARS-CoV-2 neutralizing antibodies from an individual who had recovered from COVID-19. We found that neutralizing antibodies to the NTD supersite commonly are encoded by the *IGHV1-24* gene, forming a genetic cluster that represents a public B cell clonotype. However, we also discovered a rare human antibody, COV2-3434, that recognizes a site of vulnerability on the SARS-CoV-2 S protein in the trimer interface and possesses a distinct class of functional activity. COV2-3434 disrupted the integrity of S protein trimers, inhibited cell-to-cell spread of virus in culture, and conferred protection in human ACE2 transgenic mice against SARS-CoV-2 challenge. This study provides insight about antibody targeting of the S protein trimer interface region, suggesting this region may be a site of virus vulnerability.

## INTRODUCTION

During the COVID-19 pandemic, more than 150 vaccine candidates have been developed, but only a few have been licensed. Most licensed vaccines encode the full-length spike (S) protein including two stabilizing proline mutations (S2P) of SARS-CoV-2 (Baden et al., 2021; Polack et al., 2020; Turner et al., 2021c) and have proven effective in protecting against SARS-CoV- 2 disease. Although SARS-CoV-2 vaccines have been developed at unprecedented speed, several questions remain about efficacy and the durability of protective immunity associated with serum neutralizing antibodies generated against the S protein. Efficacy studies are complicated by the emergence of SARS-CoV-2 variants of concern (VOC) that can escape some neutralizing antibodies. Antibodies that neutralize SARS-COV-2 VOC have been studied broadly by many groups, both in terms of their potency and structure (Barnes et al., 2020; Baum et al., 2020; Cerutti et al., 2021b; Chen et al., 2021b; Chi et al., 2020; Dong et al., 2021; Hansen et al., 2020; McCallum et al., 2021; Pinto et al., 2020; Rogers et al., 2020; Shi et al., 2020; Suryadevara et al., 2021; Turner et al., 2021b; Zost et al., 2020a). Similarly, it has been reported that unrelated individuals can produce genetically and functionally similar clones of antibodies (“public clonotypes”) following infection or vaccination ((Chen et al., 2021a; Robbiani et al., 2020; Soto et al., 2019; Yuan et al., 2020)*18-22*).

The receptor binding domain (RBD) of the S protein interacts with angiotensin-converting enzyme 2 (ACE2). In addition, the N-terminal domain (NTD) of S has been proposed to cooperate with receptors or co-receptors, such as dendritic cell-specific intercellular adhesion molecule-3-grabbing non-integrin (DC-SIGN or CD209), neuropilin-1 (NRP-1), and liver-/lymph-node-specific intracellular adhesion molecules-3 grabbing non-integrin (L-SIGN or CD209L) to mediate viral attachment and enable SARS-CoV-2 infection via the established ACE2 receptor pathway (Amraei et al., 2021; Cantuti-Castelvetri et al., 2020; Daly et al., 2020; Lempp et al., 2021). Furthermore, the NTD of SARS-CoV-2 spike reportedly binds biliverdin by recruitment of tetrapyrrole rings, to evade neutralization of SARS-CoV-2 by some antibodies (Rosa et al., 2021). SARS-CoV-2 S appears to exhibit conformational flexibility of divergent loop regions in the NTD to accommodate diverse glycan-rich host sialosides that may allow it to infect host cells with wide tissue tropism (Awasthi et al., 2020). Taken together, our understanding of the functional qualities of the human antibody response against NTD is incomplete. We and other groups previously have identified potently neutralizing NTD-specific mAbs targeting one major antigenic site (Cerutti et al., 2021b; Chi et al., 2020; McCallum et al., 2021; Suryadevara et al., 2021; Voss et al., 2021; Wu et al., 2020). Here, using the single-B-cell barcoding LIBRA-seq antibody discovery technology, we performed targeted discovery of NTD-reactive antibodies from an individual who had recovered from a previous SARS-CoV-2 infection. Our results indicate that a dominant human B cell response to that major NTD antigenic site comprises clones encoded by common variable gene segments (*i.e.*, constitute a “public clonotype”). The scale of antibody discovery possible with LIBRA-seq also allowed us to identify a rare clone with unusual specificity and function.

## RESULTS

### SARS-CoV-2 infection induces a strong response against NTD and durable neutralization titers

Peripheral blood samples were obtained following written informed consent from four subjects (D1988, D1989, D1995 and D1951) infected in the United States who tested PCR-positive for SARS-CoV-2 infection and one healthy donor (D269) who served as a negative control (**Table S1**). We isolated plasma or serum specimens from the five individuals and performed serum/plasma antibody ELISA binding assays using soluble proline-stabilized S ectodomain (S2P_ecto_), RBD, or NTD protein from SARS-CoV-2 or S2P_ecto_ protein from SARS-CoV. All subjects (except the negative control) had circulating antibodies that recognized each of the proteins tested, with the greatest reactivity against the SARS-CoV-2 S2P_ecto_, RBD and NTD proteins (**Fig. 1A**). The serum antibody reactivity of one individual (D1989) was highest against the SARS-CoV-2 NTD protein (**Fig. 1A**). Consequently, we focused our efforts on identifying B cells from the blood samples of this individual, using sequential collections on day 18, 28, 58 and day 90 after onset of symptoms. This individual also possessed high serum neutralizing antibody titers, as determined in an assay using a chimeric vesicular stomatitis virus (VSV) displaying SARS-CoV-2 S protein (VSV-S) in a real time cell analysis (RTCA) method (10) (**Fig. 1B**). The plasma neutralizing titer was high (NT_50_= 1:258) even three months after recovery from SARS-CoV-2 infection. To corroborate the VSV-S-based neutralization results, we also performed a serum-antibody focus reduction neutralization test (FRNT) using an authentic SARS-CoV-2 strain (WA/1/2020). The authentic virus assay gave similar results to the VSV-based assay (**Fig. 1C**).

**Figure 1.**
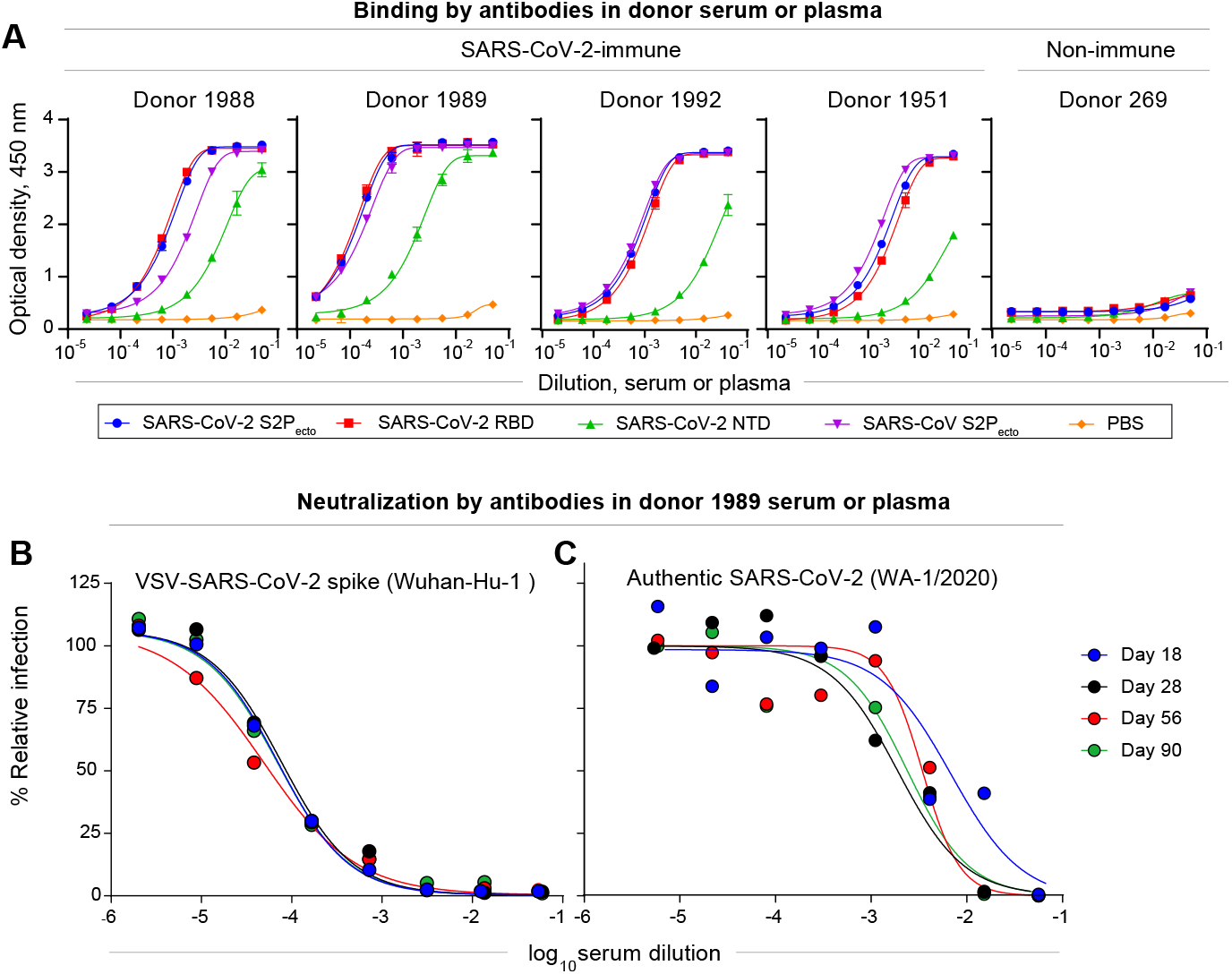
Characterization of SARS-CoV-2 antibodies in convalescent patient samples. **A.** Serum or plasma antibody reactivity for the four SARS-CoV-2 convalescent patients and one non-immune healthy control subject were assessed by ELISA using SARS-CoV-2 S6P_ecto_, S_RBD_, S_NTD_, SARS-CoV S2P_ecto_ or PBS. Optical density was measured with a 450-nm filter (OD_450_) using a microplate reader. Error bars, s.d.; data are representative of at least two independent experiments performed in technical duplicate. **B.** Plasma or serum neutralizing activity against the VSV-S for SARS-CoV-2 convalescent donor 1989 on day 18, 28, 56 or day 90 in an RTCA neutralization assay. Data represent two experiments performed in technical duplicate. **C.** Plasma or serum neutralizing activity against the WA1/2020 strain SARS-CoV-2 for convalescent donor 1989 on day 18, 28, 56 or day 90 using a FRNT. Data represent experiments performed in technical duplicate. Data represent two experiments performed in technical duplicate.

### LIBRA-seq identifies antigen-specific B cells with high NTD specificity

Next, we used the LIBRA-seq (*Linking B Cell receptor to antigen specificity through sequencing*) method (Setliff et al., 2019) to identify NTD-reactive B cells. This high-throughput technology enables determination of B cell receptor sequence and antigen reactivity at single-cell level. The LIBRA-seq antigen screening library included SARS-CoV-2 S protein stabilized in a prefusion conformation (S6P_ecto_) and NTD from SARS-CoV-2 (2019-nCoV), along with antigens from other coronavirus strains and negative control antigens. We identified 347 NTD-specific B cells from individual D1989 (day112). We recovered 108 B cells which expressed unique V_H_–JH– CDRH3–V_L_–J_L_–CDRL3 clonotypes and gave LIBRA-seq scores above a threshold of 1 for rNTD (**Fig. 2A**), and we were able to express 102 of these sequences as human mAbs. To confirm the antigen specificity predicted by the LIBRA-seq score, we tested all expressed mAbs for binding in ELISA to recombinant monomeric RBD or NTD of SARS-CoV-2 or trimeric S6P_ecto_ of SARS-CoV-2 or trimeric S2P_ecto_ of SARS-CoV proteins. We confirmed the predicted antigen specificity for greater than 90% of the clones (**Fig. 2B**). Most antibodies recognized NTD protein, except for COV2-3454 which recognized RBD (**Fig. 2B**).

**Figure 2.**
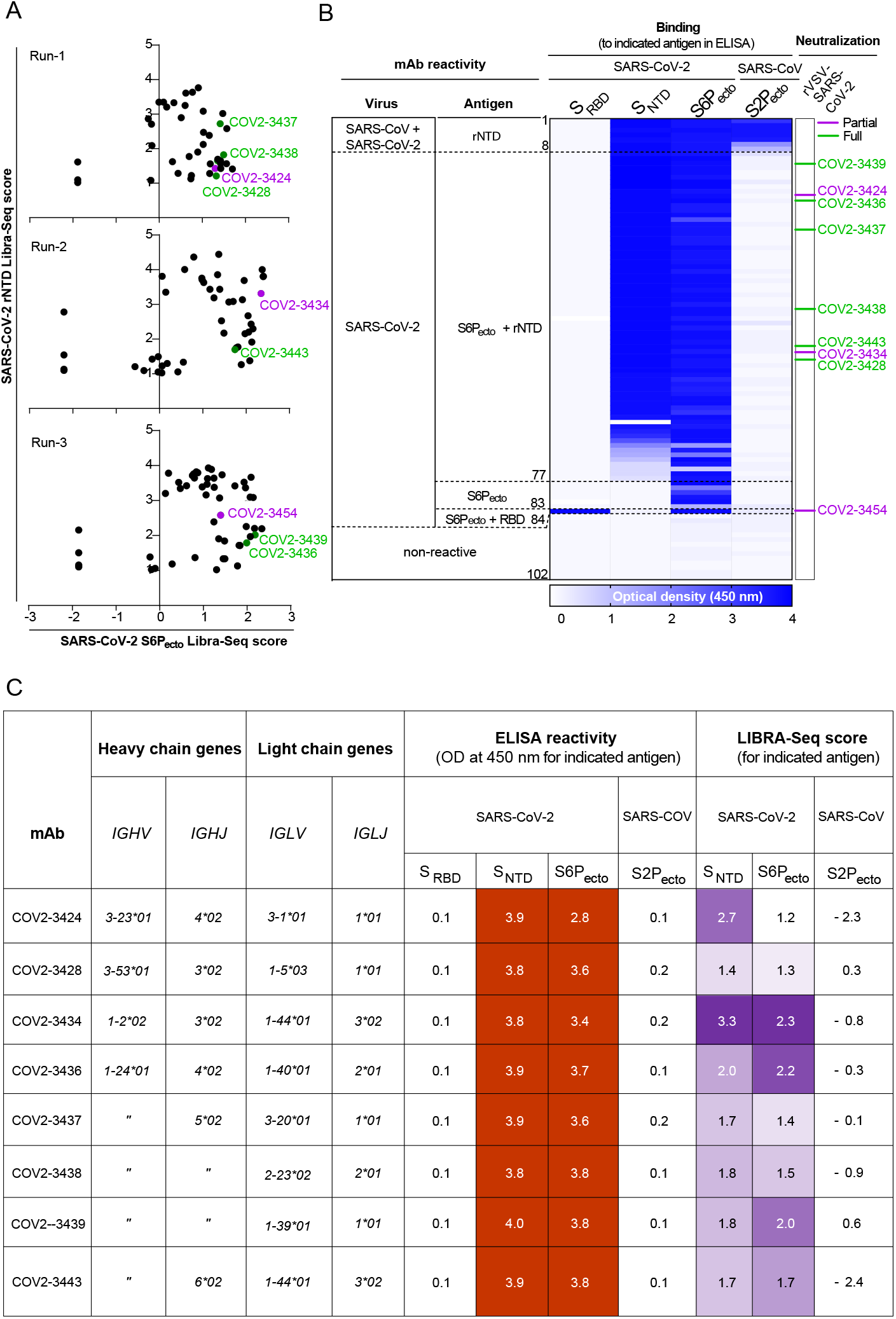
Reactivity, functional and genetic features of 102 human mAbs identified using LIBRA-seq. **A.** LIBRA seq scores for all cells per experiment are shown as black circles from three different LIBRA-seq runs. Antibodies that demonstrated either full and partial neutralization in the high-throughput RTCA assay are highlighted in green or purple, respectively. **B.** MAb specificity or reactivity for each of four S proteins or subdomains. The figure shows a heatmap for binding of 102 mAbs expressed recombinantly, representing OD values collected at nm for each antigen (range, 0.5 to 4.0). White indicates a lack of detectable binding, blue indicates binding, and darker blue indicates higher OD values. To the right are the antibody numbers that demonstrated either full or partial neutralization in the high-throughput RTCA assay, highlighted in green or purple, respectively. **C.** Genetic characteristics for mAbs that demonstrated either full or partial neutralization along with their ELISA reactivity; numbers in the boxes represent OD values collected at 450 (range, 0.5 to 4.0) and LIBRA-seq scores for each antigen. White fill indicates no or low reactivity, red (ELISA) or purple (LIBRA-seq) fill represent reactivity for the respective antigens.

Additionally, using the RTCA method, we performed high-throughput neutralization assays with VSV-S and identified 9 mAbs that showed either full (100%) or partial (50 to 80%) neutralizing capacity (**Fig. 2B**). Next, we analyzed the sequences of the variable region genes for the 102 expressed antibodies to assess the genetic diversity of antigen-specific B cell clonotypes discovered. The expressed antibodies had diverse sequence features, including varied V- and J-gene usage, CDR3 lengths, and somatic hypermutation levels for both the heavy and light chains (**Fig. S1a**). After clustering these clones based on the inferred immunoglobulin heavy variable (IGHV) gene, we noted that the *IGHV1-24* and *IGHV1-69* variable gene segments were used frequently in this individual’s response (**Fig. S1b**). Five of the nine neutralizing mAbs are encoded by the *IGHV1-24* gene segment and are clonally unrelated (**Fig. 2C**).

### Potently neutralizinge antibodies against NTD belong to public clonotypes

Next, we determined the binding activity of the panel of NTD-reactive neutralizing antibodies. Using serial dilution studies, we determined the half-maximum effective concentration (EC_50_) for binding to the S6P_ecto_ trimer protein, in comparison with a known NTD-reactive mAb (4A8) or a negative control dengue-specific antibody (rDENV-2D22). NTD-reactive neutralizing antibodies exhibited varied binding profiles with a diverse range of EC_50_ values (**Fig. 3A**). We also tested the panel of antibodies for binding to cell-surface-displayed S protein on SARS-CoV-2-infected cells according to the gating strategy shown in **Fig S2**. Unexpectedly, the NTD-targeting mAbs stained infected cells with greater intensity (higher median fluorescence intensity [MFI]) than a previously described high-affinity RBD-reactive potently neutralizing mAb (COV2-2196) (Zost et al., 2020a) (**Fig. 3B**). We also determined the inhibitory potency for representative mAbs in the quantitative VSV-S-based neutralization assay (**Fig. 3C**). These results confirmed that the LIBRA-seq technology efficiently identifies mAbs with the correct antigen specificity and that some of the NTD-reactive mAbs potently neutralize VSV-S infection based on RTCA neutralization ((Suryadevara et al., 2021; Zost et al., 2020a). Next, we chose chose COV2-3434 for further study with FRNT as it showed a distinct phenotype both in binding and rVSV neutralization. We performed FRNTs for mAb COV2-3434 using strains SARS-CoV-2 D614G and chimeric strains expressing the B.1.351 (Beta) spike in the WA1/2020 background (Wash-B 1.351) (Chen et al., 2021b). COV2-3434 neutralized both strains of SARS-CoV-2 in a dose-dependent manner, with half-maximal inhibitory (IC_50_) values of 5.5 or 32 µg/mL, respectively (**Fig. 3D**). A comprehensive analysis of antibody variable gene sequences for SARS-CoV-2 human mAbs revealed that the *IGHV1-24* gene segment is frequently used by vaccinated or convalescent individuals when targeting the NTD (**Table S2, Fig. S3**). Nevertheless, the clones recovered here were unique with diverse gene usage for both heavy and light chains. There was no bias for a particular HCDR3 length that confers NTD-specificity. Additionally, the *IGHV1-69* and *IGHV3-53* gene segments are over-represented in both RBD- and NTD-specific antibodies isolated from convalescent subjects. Of note, the *IGHV1-69* gene-encoded antibodies that reacted with NTD did *not* neutralize VSV-S, and the other V_H_ genes used (*IGHV1-2*, *IGHV3-23* and *IGHV3-53*) encoded clones with only moderate neutralizing capacity. Thus, the most potently neutralizing NTD-reactive antibodies isolated here were encoded by *IGHV1-24*.

**Figure 3.**
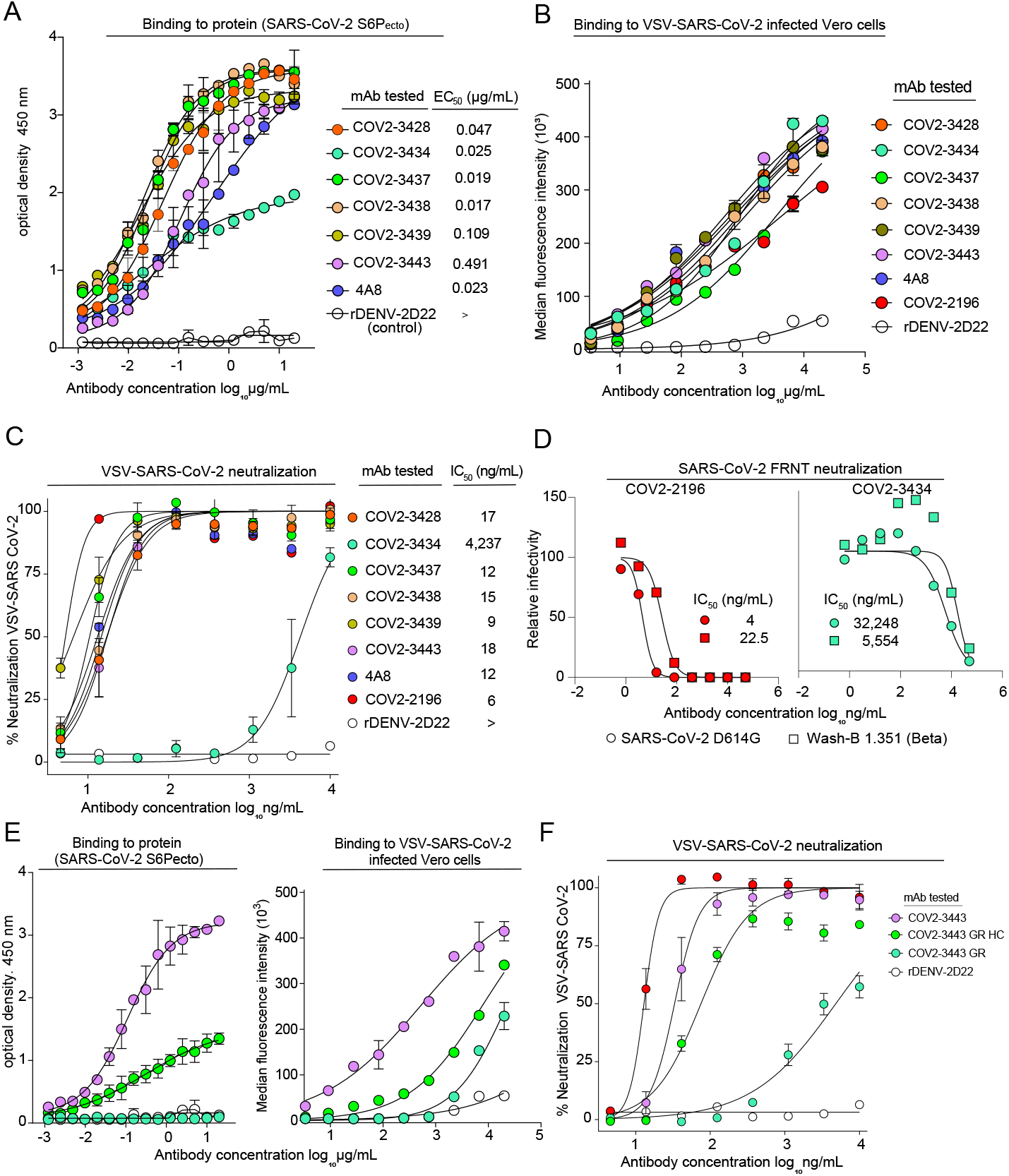
Activity of neutralizing mAbs against SARS-CoV-2. **A.** ELISA binding to SARS-CoV-2 S6P_ecto_ protein was measured by absorbance at 450 nm. Antibody concentrations starting at 20 µg/mL were used and titrated two-fold. Calculated EC_50_ values are shown on the graph. Error bars indicate standard deviation; data represent at least two independent experiments performed in technical duplicate. **B.** Binding to the surface of VSV-S-infected Vero cells was measured using flow cytometry and median florescence intensity values were determined for dose-response binding curves. Antibody was diluted 3-fold staring from 20 µg/mL. Data represent two experiments performed in technical triplicate. **C.** VSV-S neutralization curves for mAbs that were expressed after high throughput RTCA neutralization conformation. Calculated IC_50_ values are shown on the graph. Error bars indicate standard deviation; data represent at least two independent experiments performed in technical duplicate. **D.** Neutralization curves for COV2-3434 or COV2-2196 against SARS-CoV-2 virus. Calculated IC_50_ values are shown on the graph. Error bars indicate standard deviation; data represent at least two independent experiments performed in technical duplicate. **E.** Germline-revertant (GR) COV2-3443 antibody reactivity and functional activity, ELISA binding to SARS-CoV-2 S6P_ecto_ protein was measured by absorbance at 450 nm and binding to the surface of VSV-S-infected Vero cells was measured using flow cytometry and median florescence intensity values were determined for dose response binding curves. **F.** VSV-S neutralization curves for germline-revertant COV2-3443 antibody. Error bars indicate standard deviation; data represent at least two independent experiments performed in technical duplicate.

To determine if the function of *IGHV1-24*-encoded antibodies identified in this study was due to germline-encoded reactivity or the result of somatic mutations, we engineered ‘germline reversion’ (GR) recombinant antibodies that were reverted at residues that differed from the germline gene segments either in the heavy chain (GR-HC) or in both heavy and light chains (GR). After alignment of the sequences of *IGHV1-24-*encoded clones the with germline gene segment *IGHV1-24*, we chose the mAb COV2-3443 for further study, as it was the antibody with the fewest somatic mutations. We tested if the GR mAb shared similar functional properties with its somatically-mutated counterparts for binding to S protein or VSV-S neutralization. The COV2-3443 GR-HC mAb retained some binding and neutralization capacity, whereas COV2- 3443 GR completely lost binding and neutralization capacity, suggesting that the functional activities required some or all of the somatic mutations present in the matured antibody (**Fig. 3 E, F**).

### COV2-3434 maps to a distinct site from the NTD supersite

We next defined antigenic sites on the NTD by competition-binding analysis. We used SARS-CoV-2 6P_ecto_ protein to screen for NTD-reactive neutralizing mAbs that competed for binding with each other or with the previously described NTD-reactive mAbs COV2-2676 and COV2-2489 that recognize known epitope on NTD (10). We also used the previously described RBD-reactive neutralizing (COV2- 2196 and COV2-2130) or non-neutralizing (rCR3022) mAbs as controls. We identified two groups of competing mAbs in the NTD (**Fig. 4A**). The first group competed for binding to the known NTD supersite, which we and others have described previously (Cerutti et al., 2021b; Chi et al., 2020; McCallum et al., 2021; Suryadevara et al., 2021). The second competition group contains a single mAb (COV2-3434) that bound to a site distinct from the epitope of all other NTD-reactive mAbs (**Fig. 4A**). We also tested competition of COV2-3434 mAb with the recently reported antibody 5-7, which binds a hydrobphobic site on NTD. Our mAb COV2-3434 did not compete for binding with mAb 5-7 either on SARS-CoV2-6P_ecto_ or on NTD, revealing the COV2-3434 site is unique (**Fig. S4**).

**Figure 4.**
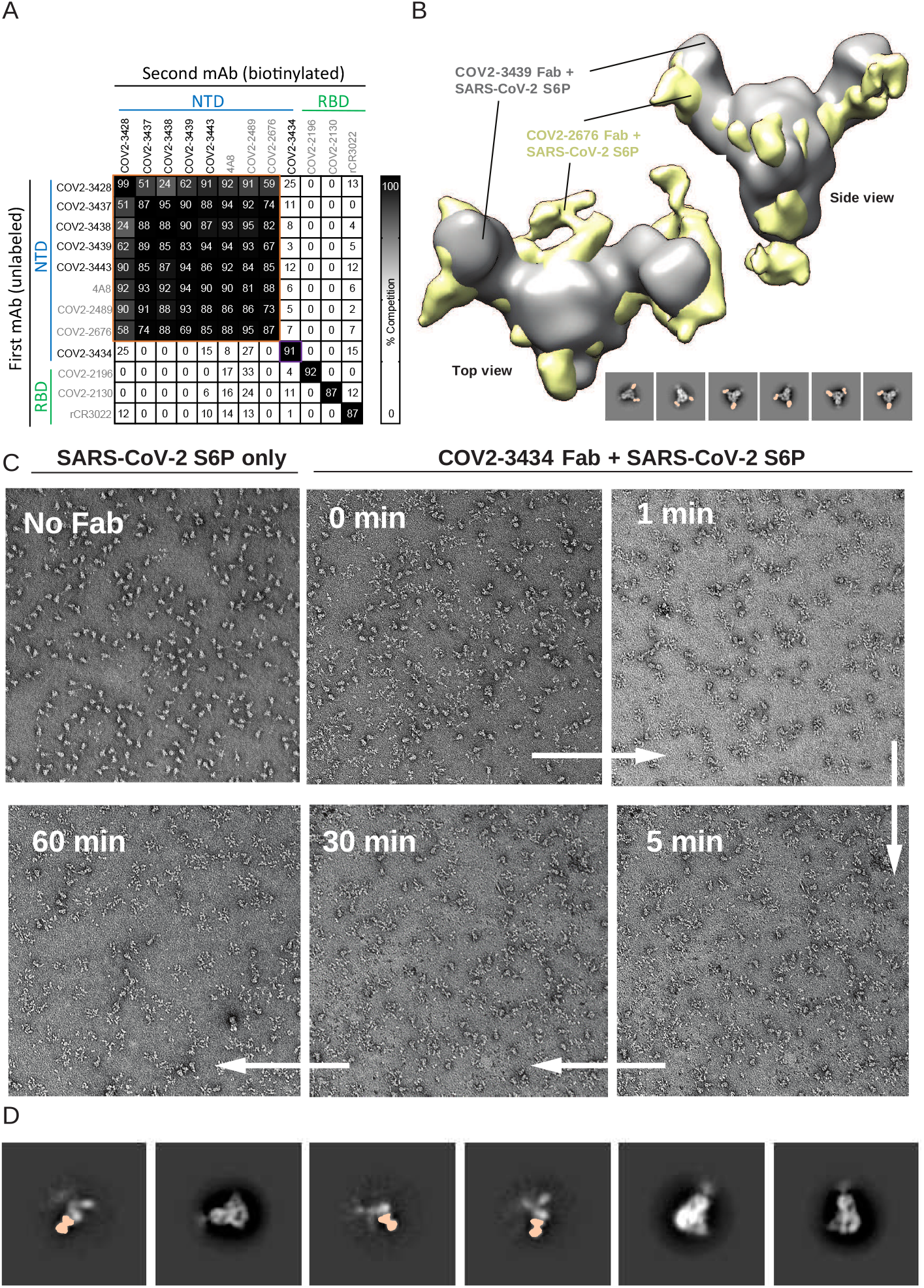
Epitope identification and structural characterization of COV2-3439 and COV2- 3434 antibodies. **A.** Competition of the panel of neutralizing mAbs with previously mapped antibodies COV2- 2130, COV2-2196, COV2-2676, COV2-2489, r4A8 or rCR3022. Unlabeled antibodies applied to antigen first are indicated on the left, while biotinylated antibodies that were added to antigen- coated wells second are listed across the top. The number in each box represents the percent competition binding of the biotinylated antibody in the presence of the indicated competing antibody. Heat map colors range from dark grey (100% binding of the biotinylated antibody) to white (0% or no binding of the biotinylated antibody). The experiment was performed in biological replicate. Biological replicate from representative single experiment shown. **B.** Negative-stain EM of SARS-CoV-2 S6P_ecto_ protein in complex with COV2-3439 Fab. Side view and top view of superimposed 3D volume COV2-3439 Fab–S6P_ecto_ closed trimer (S protein model PDB:7JJI) complexes as visualized by negative-stain EM for COV2-2676 Fab model in gold, COV2-2489 Fab model in grey. At the bottom, negative-stain 2D classes of SARS-CoV-2 S protein incubated with COV2-3439 Fab are shown. Data are from a single experiment; detailed collection statistics are provided in Supplementary Table 3. **C.** Morgagni images of SARS-CoV-2 S6P_ecto_ protein only, immediately after COV2-3434 Fab was added to SARS-CoV-2 S6P_ecto_ trimer, incubated for 1, 5, 30 mins or 1 hr and placed on an nsEM grid. **D.** Negative-stain 2D classes of SARS-CoV-2 S6P_ecto_ protein only or COV2-3434 Fab with a monomer of SARS-CoV-2 S6P_ecto_ protein (based on the density surrounding the Fab).

### COV2-3434 exhibits trimer-disrupting properties

To further probe the binding sites for these mAbs, we used negative-stain electron microscopy (nsEM) to image a stabilized trimeric form of the ectodomain of S protein (S6P_ecto_ trimer) in complex with Fab fragment forms of COV2-3439 or COV2-3434. We chose COV2-3439 as a representative mAb from the first competition group, as it was the most potently neutralizing antibody against VSV-S. COV2-3439 bound to the NTD and recognized the ‘closed’ conformational state of the S6P_ecto_ trimer. We confirmed that the COV2-3439 antibody binds to the previously noted antigenic “supersite” on the NTD of the S6P_ecto_ trimer by overlaying the nsEM maps of the COV2-3439 Fab/S protein complex with our previously published COV2-2676 Fab/S complex (**Fig. 4B**).

Unexpectedly, we did not observe intact S protein trimers following a one-hour incubation with saturating concentrations of COV2-3434 Fab fragments. Shorter incubation times with Fabs (1, 5 or 30 mins) showed more intact trimers in the grids (**Fig. 4C**). Representative 2D images revealed that Fabs were bound to the S protomers, suggesting that Fabs recognize an epitope that is not present or accessible on an intact S trimer (**Fig. 4D**). Although the 2D images are revealing, we could not create reconstructions of the Fab-protomers, since there were very limited views of the complexes. The data are consistent with a trimer-disruption mechanism in which binding of the COV2-3434 Fab to a partially occluded epitope drives the disruption of S protein trimer.

We next defined the COV2-3434 and COV2-3439 epitopes at the amino acid level using 2 complementary methods: alanine-scanning loss-of-binding experiments and cell-surface S protein display method. Screening of the NTD alanine-scan library identified primary residues F43, F175, L176 and L226 as critical for binding of COV2-3434 (**Fig. 5A**), whereas for COV2- 3439 residues R102, Y145, K147, W152, R246, Y248, P251 and G252 were identified **(Fig. S5).** None of these single-residue alanine mutants affected binding of the control NTD-reactive mAb COV2-2305 (**Fig. 5B**). As an alternative approach to learn more about the epitope recognized this trimer-disrupting antibody, we generated complexes of NTD subdomain with Fabs of COV2-3434 and COV2-3439. Intrestingly, in NS-EM we noticed that the COV2-3434 Fab binds NTD at a 90° angle to that of the supersite-binding COV2-3439 Fab (**Fig. 5C**). Moreover, when we overlaid this double Fab + rNTD complex onto that of the trimeric spike complex (7C2L model), COV2-3434 Fab tangentially clashed with interface of RBD and NTD (**Fig. 5C**). Modeling of double Fab and NTD complexes onto the spike monomer, dimer and trimer when RBD is open enabled us to locate Fab binding more precisely and suggested that the epitope recognized by COV2-3434 is occluded (**Fig. S6**). Recently, it was reported that the NTD of SARS-CoV-2 spike binds biliverdin and polysorbate 80 by recruitment of tetrapyrrole rings to evade antibody neutralization. However, our neutralization assays in the presence of biliverdin or polysorbate 80 did not affect COV2-3434 neutralization of VSV-S (**Fig. S7**), again suggesting this epitope is distinct. Additional structural studies are needed to determine sturcural basis for the trimer-disrupting phenotype of mAbs binding to this epitope.

**Figure 5.**
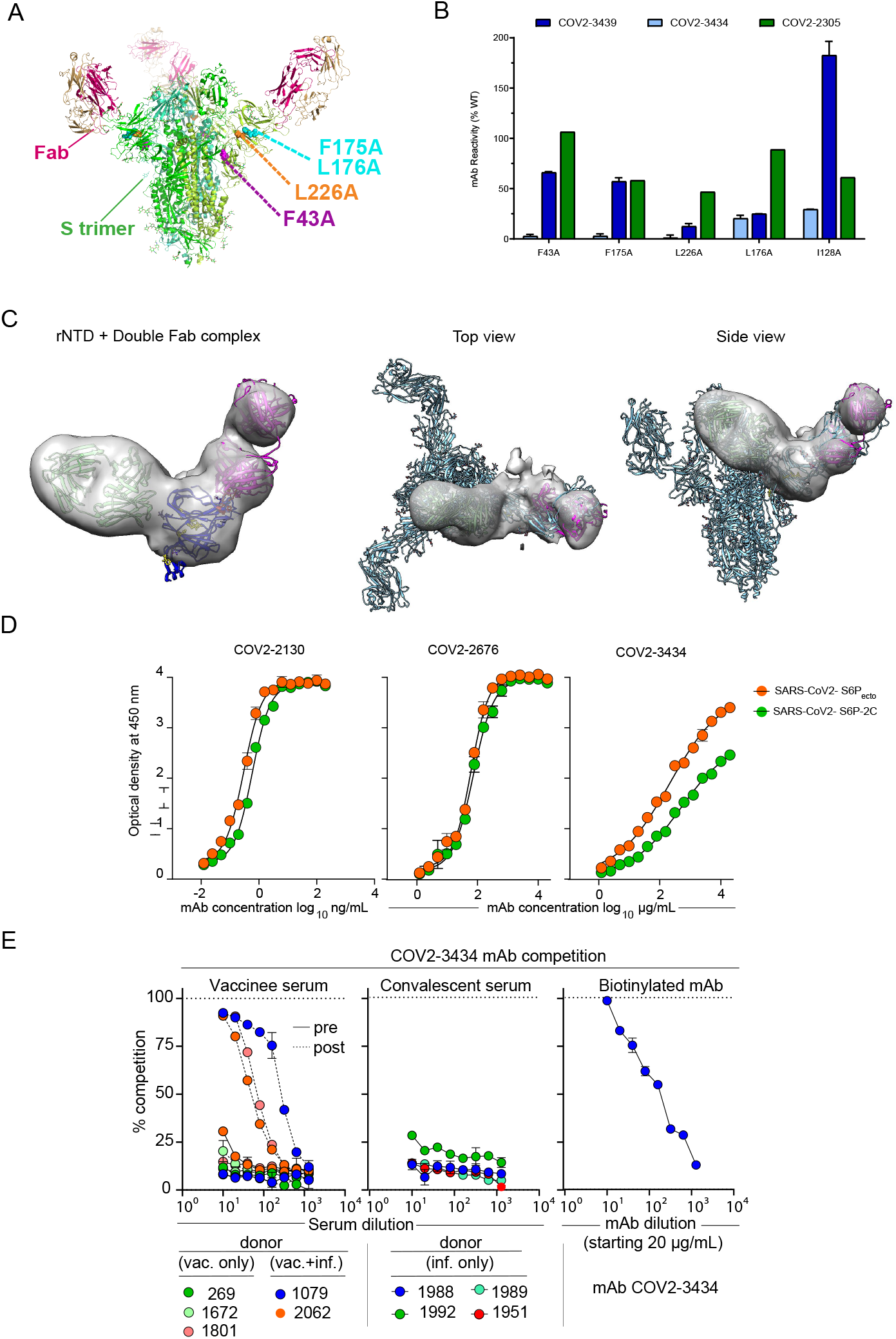
Structural characterization of the trimer-disrupting antibody COV2-3434. **A.** Residues identified as important for COV2-3434 binding are highlighted as spheres on the S protein structure (green ribbon; PDB 7L2C) F43 (magenta), F175, L176 (cyan), or L226 (orange). Residues critical for COV2-3434 binding were identified from binding screens of an alanine scanning mutagenesis library of NTD. **B.** MAb binding values for COV2-3434, COV2-3439, and control anti-NTD mAb COV2-2305 are shown at SARS-CoV-2 S protein clones identified as critical for MAb binding. MAb reactivities for each mutant are expressed as percent of binding to wild-type S protein, with ranges (half of the maximum minus minimum values). Two replicate values were obtained for each experiment. **C.** Negative-stain EM of SARS-CoV-2 rNTD protein in complex with COV2-3439 and COV2- 3434 Fabs. Top view and side view of superimposed 3D volume COV2-3434 Fab - COV2-3439 Fab - SARS-CoV-2 rNTD complexes as visualized by negative-stain EM aligned to S protein of SARS-CoV-2 in complex with 4A8 (PDB: 7C2L) Data are from a single experiment; detailed collection statistics are provided in Supplementary Table 3. **D.** ELISA binding to SARS-CoV-2 S6P_ecto_ or SARS-CoV-2 S6P-2C was measured by absorbance at 450 nm. The COV2-2130 starting concentration was 200 ng/mL, the COV2-2676 and COV2-3434 starting concentrations were 20 µg/mL, and mAbs were titrated two-fold. Calculated EC_50_ values are shown on the graph. Error bars indicate standard deviation; data represent at least two independent experiments performed in technical duplicate. **E.** Measurement of serum antibody competition with trimer interface antibody COV2-3434 in individuals before or after SARS-CoV-2 mRNA vaccination. Competition-binding ELISA curves for COV2-3434 with human serum from convalescent or vaccinated donors. Competition- binding experiments were performed for each sample in triplicate and repeated in at least 2 independent experiments. One representative experiment is shown. For all competition-binding curves, data points indicate the mean and error bars indicate the standard deviation.

The S protein exhibits high flexibility between domains and can exist in different conformations, allowing the immune system to target distinct epitopes and structural states (Henderson et al., 2020). Henderson *et al*. showed that conformations of the S protein can be controlled via rational design using expressed soluble ectodomains of the S proteins, in which the three RBDs are either locked in the all-RBDs ‘down’ position (S6P_ecto_-2C) or adopt ‘up’ state (S6P_ecto_) conformations (Henderson et al., 2020). We hypothesized that the COV2-3434 binding site is accessible only when the RBD adopts an ‘up’ state conformation of S6P_ecto_. To test this model, we quantified binding of COV2-3434 to S6P_ecto_ or S6P_ecto_-2C proteins by ELISA. For comparison, we also included a mAb that binds to RBD in either the up or down conformational state (COV2-2130), a mAb that binds to NTD (COV2-2676), and the negative-control dengue mAb DENV-r2D22. As expected, the binding of COV2-3434 to S6P_ecto_-2C protein was reduced, confirming that the epitope is cryptic and only accessible when at least one RBD is in its ‘up’ conformation (**Fig. 5D**).

### SARS-CoV-2 mRNA vaccines can induce trimer-disrupting antibodies

Although we identified a new antigenic site by isolating COV2-3434 from a SARS-CoV-2 convalescent donor, it is uncertain if this class of antibodies forms a major part of the humoral immune response to the S protein trimer. To address this question, we performed a competition-binding ELISA with serum antibody and COV2-3434. Serum antibodies from each of 4 naturally SARS-CoV-2 infected individuals or from each of 5 individuals before or after SARS-CoV-2 mRNA vaccination were tested. We observed up to 90% serum antibody competition with COV2-3434 in 3 donors tested following vaccination, indicating that in some individuals SARS-CoV-2 mRNA vaccination generates high levels of S protein trimer-interface (TI) specific antibodies or antibodies that compete with TI antibodies (**Fig. 5E**). In contrast, we did not observe this level of competition with COV2-3434 in serum from convalescent donors. Taken together, these results suggest that S trimer interface antibodies may be more common in the serum of vaccinated than infected individuals. The reason this class of antibodies was observed in the serum of vaccinees but not convalescent individuals is not clear, although engineered vaccine S antigen differs from the natural S protein in that the “pre-fusion” S conformation was stabilized in the vaccine construct by mutagenesis.

### COV2-3434 inhibits VOC and confers partial protection against SARS-CoV-2 infection

Identification of neutralizing mAbs that bind to distinct antigenic sites on S proteins might help to avoid escape from neutralization by VOC. To address this idea, we used VSV-S viruses expressing SARS-CoV-2 S protein variants that were resistant to neutralization by the RBD-specific antibodies COV2-2479, COV2-2499 or COV2-2130 (Greaney et al., 2021) or resistant to the NTD-specific antibodies COV2-2676 and COV2-2489 (Suryadevara et al., 2021). The COV2-3434 mAb neutralized all escape VSV viruses at the higher concentration tested (**Fig. 6A**).

**Figure 6.**
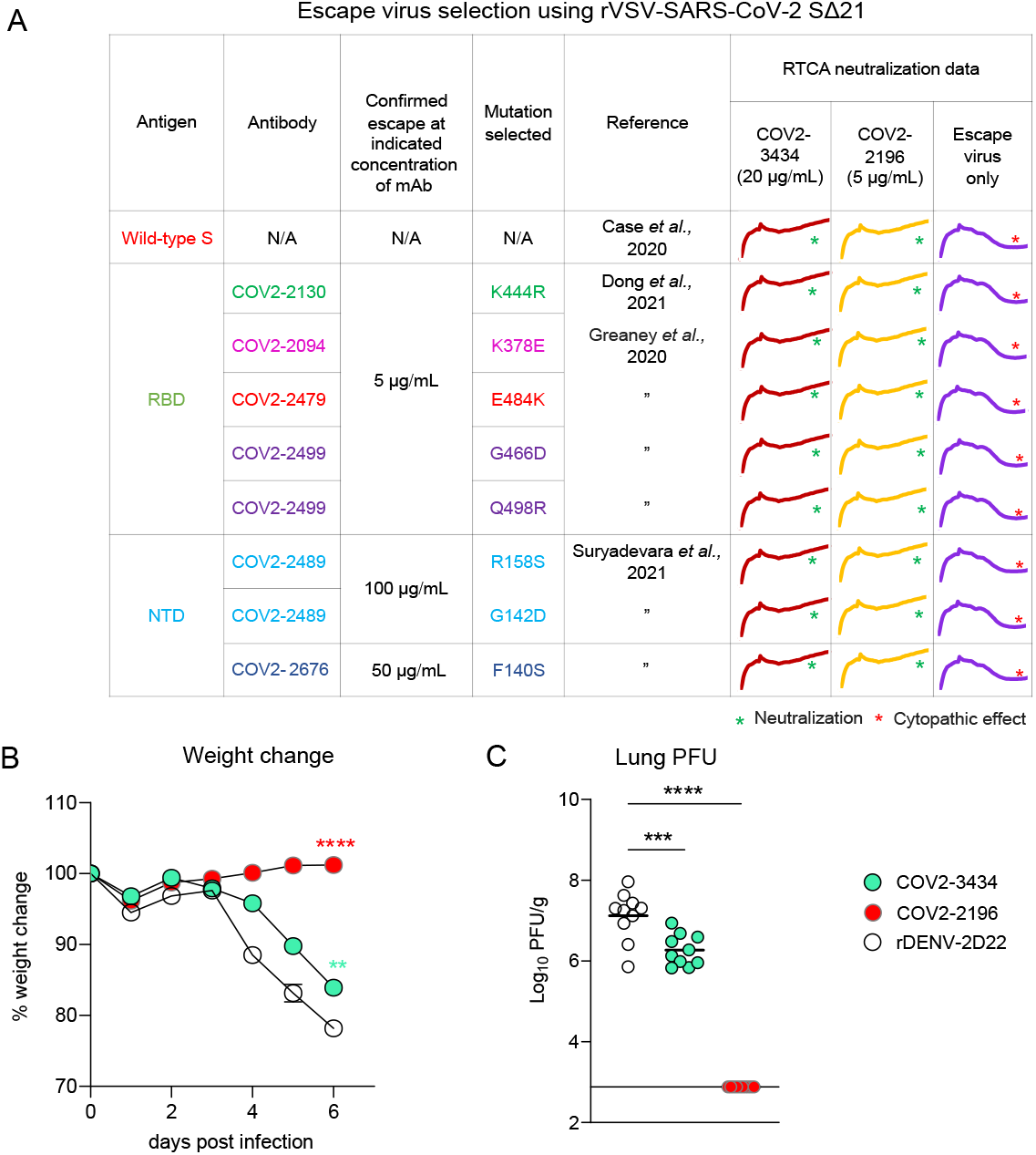
Escape virus neutralization and protection in K18 hACE2 transgenic mice by trimer-disrupting antibody COV2-3434. **A.** Neutralization of mAb escape viruses selected by RBD-specific mAbs COV2-2479 (red), COV2-2130 (green), COV2-2094 (magenta) or COV2-2499 (purple) and NTD-specific mAbs COV2-2676 (blue) or COV2-2489 (cyan) and with VSV-S by COV2-3434 or COV2-2196 (positive control). Mutations selected by those mAbs are listed with the references. Toward the right, the RTCA curves show neutralization of those escape viruses, The * symbol indicates lack of neutralization in wells with only virus and no antibody. **B.** Eight-week-old male K18-hACE2 transgenic mice were inoculated by the intranasal route with 10^4^ FFU of SARS-CoV-2 (WA1/2020 D614G). One day prior to virus inoculation, mice were given a single 200 µg (∼10 mg/kg) dose of COV2-3434 or COV2-2196 by intraperitoneal injection. Weight change was monitored daily. Data are from two independent experiments, n=10 per group. **, p<0.01; ****, p<0.0001. Error bars represent SEM. **C.** At 6 dpi, lungs were collected and assessed for infectious viral burden by plaque assay. Plaque-forming units (PFU)/g is shown. Bars indicate the mean viral load; the dotted line indicates the limit of detection of the assay. Data are from two independent experiments, n=10 per group. **, p<0.01; ****, p<0.0001.

We next assessed the ability of COV2-3434 to protect K18-hACE2-transgenic mice following viral challenge with SARS-CoV-2 (Golden et al., 2020; Oladunni et al., 2020; Winkler et al., 2021). One day prior to virus inoculation, we passively transferred ∼10 mg/kg (200 µg/mouse) of COV2-2196 (RBD-specific), COV2-3434 (NTD-specific) or DENV-r2D22 (negative control) mAbs. Mice that received r2D22 lost more than 20% initial body weight. Animals treated with the RBD mAb COV2-2196 were completely protected from weight loss. COV2-3434 conferred intermediate protection against weight loss (**Fig. 6B**). Pre-treatment with COV2-3434 also partially protected against viral burden, with a 7-fold lower level of infectious virus in the lung compared to the negative-control antibody (**Fig 6C**). We repeated the study by passively transferring a higher dose (1 mg/mouse) of COV2-2196 (RBD-specific), COV2-3434 (NTD-specific) or DENV-r2D22 (negative control) mAbs, and again saw a comparable reduction of viral titers in the lungs and nasal turbinates (**Fig. S8**).

## DISCUSSION

Human neutralizing mAbs to SARS-CoV-2 isolated from recovered COVID-19 individuals are of great importance as potential therapeutic candidates. The continued investigation into identifying protective epitopes using mAbs as we have done here may inform future structure-based rational design of next-generation SARS-CoV-2 vaccines by revealing protective sites whose structure should be preserved in engineered vaccine antigens. Most potently neutralizing SARS-CoV-2 mAbs discovered to date recognize the RBD region, while some moderately neutralizing NTD-directed mAbs also were identified (Barnes et al., 2020; Baum et al., 2020; Cerutti et al., 2021b; Chen et al., 2021b; Chi et al., 2020; Dong et al., 2021; Hansen et al., 2020; McCallum et al., 2021; Pinto et al., 2020; Rogers et al., 2020; Shi et al., 2020; Suryadevara et al., 2021; Turner et al., 2021c; Zost et al., 2020a). All of the NTD-reactive mAbs reported to date have lost their neutralizing capacity against certain emerging VOC. The majority of antibodies identified against NTD target an antigenic site termed the NTD ‘supersite’ (Cerutti et al., 2021b; Chi et al., 2020; McCallum et al., 2021; Suryadevara et al., 2021). Although a few other antigenic sites on NTD have been described, mAbs binding to these sites generally were non-neutralizing. The frequent occurrence of mutations in the NTD of multiple circulating SARS-CoV-2 variants suggests that the NTD is under strong selective pressure from the host humoral immune response (Weisblum et al., 2020). Furthermore, antigenic changes caused by deletions in NTD have been identified within the antigenic supersite of viruses shed by immunocompromised hosts (Avanzato et al., 2020; Choi et al., 2020; McCarthy et al., 2021).

In this work, we report the isolation and characterization of SARS-CoV-2 neutralizing mAbs targeting the NTD using LIBRA-seq. We used NTD, a domain cloned from the full-length spike, as antigen bait for isolating memory B cells from a convalescent donor. More than 90% of the clones we selected by LIBRA-seq for expression reacted exclusively with NTD, and these findings also were supported by reactivity studies with the SARS-CoV-2 S6P_ecto_ domain. A subset of eight NTD-targeting antibodies selected by LIBRA-seq was neutralizing. Several of the mAbs potently neutralized VSV-S. The primary target for most of the neutralizing antibodies identified is the NTD ‘supersite’, as previously described by several groups (Cerutti et al., 2021b; McCallum et al., 2021; Shi et al., 2020; Suryadevara et al., 2021). Most of these NTD-supersite-targeting antibodies appear to be members of a public clonotype. Although diverse public clonotypes recognizing RBD or NTD have been described, we identified an *IGHV1-24*- encoded clonotype that seems to dominate the response to NTD. Clones from this public clonotype are seen following both vaccination and infection.

We also identified an antibody designated COV2-3434 that recognizes a distinct antigenic site on NTD that may represent a new site of vulnerability on SARS-CoV-2 spike. COV2-3434 binds to recombinant SARS-CoV-2 S6P_ecto_ protein weakly in ELISA, but more avidly to cell-surface-displayed spike on Vero cells infected with VSV-S. In contrast to other NTD-reactive potently neutralizing antibodies, COV2-3434 weakly inhibits infection of VSV-S and authentic SARS-CoV-2 viruses. With these distinctive phenotypes, we tried to learn more about the mode of recognition of this antigenic site by ns-EM of antigen-antibody complexes. Unexpectedly, we found that COV2-3434 Fab disrupted SARS-CoV-2 S trimers when added to make spike-Fab complexes. This finding of trimer disassociation mediated by COV2-3434 revealed a potential site of vulnerability hidden in the S trimer interface. Similarly, a recently identified NTD-reactive neutralizing antibody called 5-7 also recognizes a distinct antigenic site within the NTD, antibodies of this class insert an antibody hypervariable loop into the exposed hydrophobic pocket between the two sheets of the NTD β-sandwich (Cerutti et al., 2021a). This pocket was described previously as the binding site for metabolites such as heme with hydrophobic properties (Rosa et al., 2021). Our alanine scan mutagenesis data reveals that COV2-3434 shares some contact residues with mAb 5-7 including F175 and L176, while L226 is barely deeper than 175 and 176. However, COV2-3434 also lost its binding capacity when deeper pocket residues F43 and was mutated. We noted that in the spike trimer, residue F43 lies at an interface between adjacent monomers such that MAb binding could intiate a destabilization of the trimer.

Recently several reports about mAbs targeting the trimer interface of multiple viral antigens have been published. For instance, the non-neutralizing influenza mAbs FluA20 and 5J6 that recognize the hemagglutinin trimer interface (Bangaru et al., 2019; Zost et al., 2021) were identified from influenza-vaccinated individuals. Also, epitope mapping using polyclonal serum from vaccinated rabbits identified antibodies recognizing the HIV envelope glycoprotein trimer interface (Turner et al., 2021a). Similarly, the epitope for a neutralizing mAb for human metapneumovirus (MPV458) lies within the trimeric interface of pneumovirus fusion proteins (Huang et al., 2020).

COV2-3434 is a rare SARS-CoV-2 S trimer interface antibody that mediates virus neutralization. Our COV2-3434 competition data suggest that this class of mAbs may be common in the serum of some vaccinated individuals. Hence, surveillance of this class of antibodies and understanding its contribution to vaccine protection is important, particularly in the context of emergence of new VOC and updated vaccine designs. While these trimer-interface mAbs do not all neutralize virus *in vitro*, passive transfer of these mAbs can mitigate severe disease. For example, the FluA20 mAb did not neutralize influenza, but still conferred protection in mice challenged with H1N1 A/California/04/2009 virus (Bangaru et al., 2019). Here, the moderately neutralizing COV2-3434 conferred partial protection against weight loss and lung infection in mice when given as prophylaxis.

In summary, using LIBRA-seq, we identified the mAb COV2-3434 that binds to a distinct antigenic site on the NTD and disassociates S trimers by contacting critical residues in a cryptic hydrophobic pocket in the S trimer interface.

## ACKNOWLEDGEMENTS

We thank Merissa Mayo and Norma Suazo Galeano for human subject’s regulatory support. EM data collection was conducted at the Center for Structural Biology Cryo-EM Facility at Vanderbilt University. This work was supported by the NIAID/NIH grants R01 AI157155 (M.S.D. and J.E.C.), R01 AI131722-S1 (I.S.G.), HHSN contracts 75N93019C00074 (J.E.C.) and 75N93019C00073 (B.J.D.), DARPA grant HR0011-18-2-0001 (J.E.C.), the Dolly Parton COVID-19 Research Fund at Vanderbilt (J.E.C.), Hays Foundation COVID-19 Research Fund (I.S.G.), and Fast Grants, Mercatus Center, George Mason University (J.E.C. and I.S.G.). J.E.C. is a recipient of the 2019 Future Insight Prize from Merck KGaA. J.B.C. is supported by a Helen Hay Whitney Foundation postdoctoral fellowship. We thank Dr. Jason McLellan for a gift of S6P_ecto_ protein used in the LIBRAseq studies. Recombinant SARS-CoV-2 S NTD protein was kindly provided by P. McTamney, K. Ren and A. Barnes (AstraZeneca). The content is solely the responsibility of the authors and does not represent the official views of the U.S. government or other sponsors.

## AUTHOR CONTRIBUTIONS

Conceived of the project: N.S., J.E.C.; Obtained funding: M.S.D., I.S.G and J.E.C. Performed laboratory experiments: N.S., A.S., R.E.C., E.B., L.V.B., J.B.C., K.K., L.M., A.T., S.M.D., L.S.H., R.N., C.N.S., E.D., Supervised research: B.J.D., I.G., R.H.C., J.E.C. Wrote the first draft of the paper: N.S., J.E.C.; All authors reviewed and approved the final manuscript.

## STAR METHODS

### RESOURCE AVAILABILITY

#### LEAD CONTACT

Further information and requests for resources and reagents should be directed to and will be fulfilled by the Lead Contact, James E. Crowe, Jr.

#### MATERIALS AVAILABILITY

Materials described in this paper are available for distribution for nonprofit use using templated documents from Association of University Technology Managers “Toolkit MTAs”, available at: https://autm.net/surveys-and-tools/agreements/material-transfer-agreements/mta-toolkit.

#### DATA AND CODE AVAILABILITY

All data needed to evaluate the conclusions in the paper are present in the paper or the Supplemental Information. The antibodies in this study are available by Material Transfer Agreement with Vanderbilt University Medical Center.

### EXPERIMENTAL MODEL AND SUBJECT DETAILS

#### Research participants

We studied the peripheral blood B cells from four individuals with a history of laboratory-confirmed symptomatic SARS-CoV-2 infection. The study was approved by the Institutional Review Board of Vanderbilt University Medical Center and specimens were obtained after written informed consent.

#### Cell lines

Vero (ATCC, CCL-81), HEK293 (ATCC, CRL-1573) and HEK293T (ATCC, CRL-3216) cells were maintained at 37°C in 5% CO2 in Dulbecco’s minimal essential medium (DMEM) containing 10% (v/v) heat-inactivated fetal bovine serum (FBS), 10 mM HEPES pH 7.3, 1 mM sodium pyruvate, 1× non-essential amino acids and 100 U/mL of penicillin–streptomycin. Vero-furin cells were obtained from T. Pierson (NIAID, NIH) and have been described previously (45) Vero-hACE2-TMPRSS2 cells were a gift of A. Creanga and B. Graham (Vaccine Research Center, NIH). FreeStyle 293F cells (Thermo Fisher Scientific, R79007) were maintained at 37°C in 8% CO2. Expi293F cells (Thermo Fisher Scientific, A1452) were maintained at 37°C in 8% CO2 in Expi293F Expression Medium (Thermo Fisher Scientific, A1435102). ExpiCHO cells (Thermo Fisher Scientific, A29127) were maintained at 37°C in 8% CO2 in ExpiCHO Expression Medium (Thermo Fisher Scientific, A2910002). Mycoplasma testing of Expi293F and ExpiCHO cultures was performed monthly using a PCR- based mycoplasma detection kit (ATCC, 30-1012K).

#### Antigen purification

A variety of recombinant soluble protein antigens were used in the LIBRA-seq experiment and other experimental assays. For the LIBRA-seq experiment, we used the S6Pecto construct. This plasmid encoded residues 1–1,208 of the SARS-CoV-2 S protein with a mutated S1/S2 cleavage site, proline substitutions at positions 817, 892, 899, 942, 986 and 987, and a C-terminal T4-fibritin trimerization motif, an 8x HisTag, and a TwinStrepTag (SARS- CoV-2 spike HP). DNA encoding this construct was transiently transfected with PEI in Expi293F cells and after six days of expression, supernatants were harvested, and protein was affinity-purified over a StrepTrap HP column (Cytiva Life Sciences). Protein was further resolved to homogeneity over a Superose 6 Increase column (GE Life Sciences).

We generated a plasmid containing a synthesized cDNA encoding a protein designate SARS - CoV-2 S-2P that possessed residues 1–1,208 of the SARS-CoV-2 spike protein as described ((Wrapp et al., 2020)46) with a mutated S1/S2 cleavage site, proline substitutions at amino acid positions 986 and 987, a C-terminal T4-fibritin trimerization motif, an 8x HisTag, and a TwinStrepTag. The plasmids were transiently transfected into FreeStyle 293F cells (Thermo Fisher Scientific) using polyethylenimine. The design of the two-proline (2P) forms of the coronavirus trimer spike antigens results in a prefusion-stabilized conformation that better represents neutralization-sensitive epitopes in comparison to their wild-type forms. Two h after transfection, cells were treated with kifunensine to ensure uniform glycosylation. Transfected supernatants were harvested after 6 days of expression.

SARS-CoV-2 S1 (cat. no: 40591-V08B1), SARS-CoV-2 S2 (cat. no: 40590-V08B), SARS-CoV-2 RBD (cat. no: 40592-V05H) and SARS-CoV-2 NTD (cat. no: 40591-V41H-B-20) truncated proteins were purchased (Sino Biological).

A gene encoding the ectodomain of a pre-fusion conformation-stabilized SARS-CoV-2 S protein ectodomain (S6Pecto) (Hsieh et al., 2020) was synthesized and cloned into a DNA plasmid expression vector for mammalian cells. A similarly designed S protein antigen with two prolines and removal of the furin cleavage site for stabilization of the prefusion form of S (S2Pecto) was reported previously (Wrapp et al., 2020). In brief, this gene includes the ectodomain of SARS-CoV-2 (to residue 1,208), a T4 fibritin trimerization domain, an AviTag site-specific biotinylation sequence and a C-terminal 8× His tag. To stabilize the construct in the pre-fusion conformation, we included substitutions F817P, A892P, A899P, A942P, K986P and V987P and mutated the furin cleavage site at residues 682–685 from RRAR to ASVG. The recombinant S6Pecto protein was isolated by metal affinity chromatography on HisTrap Excel columns (Cytiva), and protein preparations were purified further by size-exclusion chromatography on a Superose 6 Increase 10/300 column (Cytiva). The presence of trimeric, pre-fusion conformation S protein was verified by negative-stain electron microscopy (Zost et al., 2020b). For electron microscopy with S protein and Fabs, we expressed a variant of S6Pecto lacking an AviTag but containing a C-terminal Twin-Strep-tag, similar to that described previously (Zost et al., 2020b). Expressed protein was isolated by metal affinity chromatography on HisTrap Excel columns (Cytiva), followed by further purification on a StrepTrap HP column (Cytiva) and size-exclusion chromatography on TSKgel G4000SWXL (TOSOH).

#### Mouse models

Animal studies were carried out in accordance with the recommendations in the Guide for the Care and Use of Laboratory Animals of the National Institutes of Health. The protocols were approved by the Institutional Animal Care and Use Committee at the Washington University School of Medicine (assurance number A3381–01). Virus inoculations were performed under anesthesia that was induced and maintained with ketamine hydrochloride and xylazine, and all efforts were made to minimize animal suffering. Heterozygous K18-hACE c57BL/6J mice (strain: 2B6.Cg-Tg(K18-ACE2)2Prlmn/J) were obtained from Jackson Laboratory (034860). Eight to nine week-old mice of both sexes were inoculated with 103 PFU of SARS-CoV-2 by an intranasal route.

#### DNA-barcoding of antigens

We used oligonucleotides that possess a 15-basepair antigen barcode, a sequence capable of annealing to the template-switch oligonucleotide that is part of the 10X Genomics bead-delivered oligonucleotides and contain truncated TruSeq small RNA read-1 sequences in the following structure: 5’-CCTTGGCACCCGAGAATTCCANNNNNNNNNNNNNCCCATATAAGA*A*A-3’, where Ns represent the antigen barcode as previously described (Setliff et al., 2019). For each antigen, a unique DNA barcode was directly conjugated to the antigen itself. In particular, 5’amino-oligonucleotides were conjugated directly to each antigen using the SoluLINK Protein-Oligonucleotide Conjugation Kit (TriLink cat. no. S-9011) according to manufacturer’s instructions. Briefly, the oligonucleotide and protein were desalted, and then the amino-oligo was modified with the 4FB crosslinker, and the biotinylated antigen protein was modified with S-HyNic. Then, the 4FB-oligo and the HyNic-antigen were mixed. This action causes a stable bond to form between the protein and the oligonucleotide. The concentration of the antigen-oligo conjugates was determined by a BCA assay, and the HyNic molar substitution ratio of the antigen-oligo conjugates was analyzed using a NanoDrop instrument according to the SoluLINK protocol guidelines. Chromatography separation on an AKTA FPLC instrument was used to remove excess oligonucleotide from the protein-oligo conjugates, which were also verified using SDS-PAGE with a silver stain. Antigen-oligo conjugates also were used in flow cytometry titration experiments.

### METHOD DETAILS

#### Antigen-specific B cell sorting

Cells were stained and mixed with DNA-barcoded antigens and other antibodies, and then sorted using fluorescence activated cell sorting (FACS). First, cells were counted, and viability was assessed using Trypan Blue. Then, cells were washed three times with DPBS supplemented with 0.1% bovine serum albumin (BSA). Cells were resuspended in DPBS-BSA and stained with cell markers including viability dye (Ghost Red 780), CD14-APC-Cy7, CD3-FITC, CD19-BV711, and IgG-PE-Cy5. Additionally, antigen-oligo conjugates were added to the stain. After staining in the dark for 30 min at room temperature, cells were washed three times with DPBS-BSA at 300 x g for five min. Cells then were incubated for 15 min at room temperature with Streptavidin-PE to label cells with bound antigen. Cells were washed three times with DPBS-BSA, resuspended in DPBS, and sorted by FACS. Antigen-positive cells were bulk sorted and delivered to the Vanderbilt Technologies for Advanced Genomics (VANTAGE) sequencing core laboratory at an appropriate target concentration for 10X Genomics library preparation and subsequent sequence analysis. FACS data were analyzed using FlowJo™ Software (Mac) version 10.6 (Becton, Dickinson).

#### Sample preparation, library preparation, and sequencing

Single-cell suspensions were loaded onto a Chromium Controller microfluidics device (10X Genomics) and processed using the B-cell Single Cell V(D)J solution according to manufacturer’s suggestions for a target capture of 10,000 B cells per 1/8 10X cassette, with minor modifications to intercept, amplify and purify the antigen barcode libraries as previously described (Setliff et al., 2019).

#### Sequence processing and bioinformatic analysis

We used our previously described pipeline to use paired-end FASTQ files of oligo libraries as input, process and annotate reads for cell barcode, UMI, and antigen barcode, and generate a cell barcode - antigen barcode UMI count matrix ((Setliff et al., 2019; Shiakolas et al., 2021). BCR contigs were processed using Cell Ranger software (10X Genomics) using GRCh38 as reference. Antigen barcode libraries were also processed using Cell Ranger. The overlapping cell barcodes between the two libraries were used as the basis of the subsequent analysis. We removed cell barcodes that had only non-functional heavy chain sequences and cells with multiple functional heavy chain sequences and/or multiple functional light chain sequences, reasoning that these may be multiplets. Additionally, we aligned the BCR contigs (filtered_contigs.fasta file output by Cell Ranger, 10X Genomics) to IMGT reference genes using HighV-Quest (Alamyar et al., 2012). The output of HighV-Quest was parsed using ChangeO (Gupta et al., 2015) and merged with an antigen barcode UMI count matrix. Finally, we determined the LIBRA-seq score for each antigen in the library for every cell as previously described (Setliff et al., 2019).

#### High-throughput antibody expression

For high-throughput production of recombinant antibodies, approaches were used that are designated as microscale. For antibody expression, microscale transfections were performed (∼1 mL per antibody) of Chinese hamster ovary (CHO) cell cultures using the Gibco ExpiCHO Expression System and a protocol for deep 96- well blocks (Thermo Fisher Scientific). In brief, synthesized antibody-encoding DNA (∼2 μg per transfection) was added to OptiPro serum free medium (OptiPro SFM), incubated with ExpiFectamine CHO Reagent and added to 800 µL of ExpiCHO cell cultures into 96-deep-well blocks using a ViaFlo 384 liquid handler (Integra Biosciences). The plates were incubated on an orbital shaker at 1,000 r.p.m. with an orbital diameter of 3 mm at 37°C in 8% CO2. The day after transfection, ExpiFectamine CHO Enhancer and ExpiCHO Feed reagents (Thermo Fisher Scientific) were added to the cells, followed by 4d incubation for a total of 5d at 37°C in 8% CO2. Culture supernatants were collected after centrifuging the blocks at 450 x g for 5 min and were stored at 4°C until use. For high-throughput microscale antibody purification, fritted deep-well plates were used containing 25 μL of settled protein G resin (GE Healthcare Life Sciences) per well. Clarified culture supernatants were incubated with protein G resin for antibody capturing, washed with PBS using a 96-well plate manifold base (Qiagen) connected to the vacuum and eluted into 96-well PCR plates using 86 μL of 0.1M glycine-HCL buffer pH 2.7. After neutralization with 14 μL of 1 M Tris-HCl pH 8.0, purified antibodies were buffer-exchanged into PBS using Zeba Spin Desalting Plates (Thermo Fisher Scientific) and stored at 4°C until use.

#### MAb production and purification

cDNAs encoding mAbs of interest were synthesized (Twist Bioscience) and cloned into an IgG1 monocistronic expression vector (designated as pTwist-mCis_G1) or Fab expression vector (designated as pTwist-mCis_FAB) and used for production in mammalian cell culture. This vector contains an enhanced 2A sequence and GSG linker that allows for the simultaneous expression of mAb heavy and light chain genes from a single construct upon transfection (Chng et al., 2015). For antibody production, we performed transfection of ExpiCHO cell cultures using the Gibco ExpiCHO Expression System as described by the vendor. IgG molecules were purified from culture supernatants using HiTrap MabSelect SuRe (Cytiva) on a 24-column parallel protein chromatography system (Protein BioSolutions).

Fab proteins were purified using CaptureSelect column (Thermo Fisher Scientific). Purified antibodies were buffer-exchanged into PBS, concentrated using Amicon Ultra-4 50-kDa (IgG) or 30 kDa (Fab) centrifugal filter units (Millipore Sigma) and stored at 4°C until use. F(ab¢)2 fragments were generated after cleavage of IgG with IdeS protease (Promega) and then purified using TALON metal affinity resin (Takara) to remove the enzyme and protein A agarose (Pierce) to remove the Fc fragment. Purified mAbs were tested routinely for endotoxin levels and found to be less than 30 EU per mg IgG. Endotoxin testing was performed using the PTS201F cartridge (Charles River), with a sensitivity range from 10 to 0.1 EU per mL, and an Endosafe Nexgen- MCS instrument (Charles River).

#### ELISA binding assays

Wells of 96-well microtiter plates were coated with purified recombinant SARS-CoV-2 S6Pecto, SARS-CoV-2 S NTD, or SARS-CoV-2 RBS protein at 4°C overnight. Plates were blocked with 2% non-fat dry milk and 2% normal goat serum in Dulbecco’s phosphate-buffered saline (DPBS) containing 0.05% Tween-20 (DPBS-T) for 1 h. The bound antibodies were detected using goat anti-human IgG conjugated with horseradish peroxidase (HRP) (Southern Biotech, cat. 2040-05, lot B3919-XD29, 1:5,000 dilution) and a 3,3′,5,5′-tetramethylbenzidine (TMB) substrate (Thermo Fisher Scientific). Color development was monitored, 1 M HCl was added to stop the reaction, and the absorbance was measured at 450 nm using a spectrophotometer (Biotek). For dose–response assays, serial dilutions of purified mAbs were applied to the wells in triplicate, and antibody binding was detected as detailed above. Half maximal effective concentration (EC_50_) values for binding were determined using Prism v.8.0 software (GraphPad) after log transformation of the mAb concentration using sigmoidal dose–response nonlinear regression analysis.

#### Cell-surface antigen-display assay

Vero cell monolayers were monitored until 80% confluent and then inoculated with VSV-SARS-CoV-2 virus (Wa1/2020 strain) (designated here as VSV-S) at an MOI of 0.5 in culture medium (DMEM with 2% FBS). For a T-225 flask, 10 mL of diluted VSV-S virus was added to the monolayer, then incubated for 40 min. During the incubation, the flask was gently rocked back and forth every 10 min to ensure even infection. Following, the incubation the flask volume was topped off to 30 mL with 2% FBS containing DMEM and incubated for 14 h. Cells were monitored for CPE under a microscope, were trypsinized and washed in FACS buffer. 100,000 infected cells were seeded per well to stain with respective antibodies. All antibody was diluted to 10 µg/mL in FACS buffer, and then serially diluted 3-fold 7 times to stain for antibodies that react to cell-surface-displayed S protein. Infected cells then were resuspended in 50 µL of diluted antibody. Antibody binding was detected with anti-IgG Alexa-Fluor-647-labelled secondary antibodies. Cells were analyzed on an iQue cytometer for staining first by gating to identify infected cells as indicated by GFP-positive cells, and then gated for secondary antibody binding.

#### Focus reduction neutralization test (FRNT)

Serial dilutions of serum/plasma were incubated with 102 FFU of SARS-CoV-2 for 1 h at 37°C. The antibody-virus complexes were added to Vero E6 cell-culture monolayers in 96-well plates for 1 h at 37°C. Cells then were overlaid with 1% (w/v) methylcellulose in minimum essential medium (MEM) supplemented to contain 2% heat-inactivated FBS. Plates were fixed 30 h later by removing overlays and fixed with 4% paraformaldehyde (PFA) in PBS for 20 min at room temperature. The plates were incubated sequentially with 1 μg/mL of rCR3022 anti-S antibody or a murine anti-SARS-CoV-2 mAb, SARS2-16 (hybridoma supernatant diluted 1:6,000 to a final concentration of ∼20 ng/mL) and then HRP-conjugated goat anti-human IgG (Sigma-Aldrich, A6029) in PBS supplemented with 0.1% (w/v) saponin (Sigma) and 0.1% BSA. SARS-CoV-2-infected cell foci were visualized using TrueBlue peroxidase substrate (KPL) and quantitated on an ImmunoSpot 5.0.37 Macro Analyzer (Cellular Technologies). Half maximal inhibitory concentration (IC_50_) values were determined by nonlinear regression analysis (with a variable slope) using Prism software.

#### High-throughput real-time cell analysis (RTCA) neutralization assay

To screen for neutralizing activity in the panel of recombinantly expressed mAbs, we used a high-throughput and quantitative RTCA assay and xCelligence RTCA HT Analyzer (ACEA Biosciences) that assesses kinetic changes in cell physiology, including virus-induced cytopathic effect (CPE). Twenty µL of cell culture medium (DMEM supplemented with 2% FBS) was added to each well of a 384-well E-plate using a ViaFlo384 liquid handler (Integra Biosciences) to obtain background reading. Six thousand (6,000) Vero-furin cells in 20 μL of cell culture medium were seeded per well, and the plate was placed on the analyzer. Sensograms were visualized using RTCA HT software version 1.0.1 (ACEA Biosciences). For a screening neutralization assay, equal amounts of virus were mixed with micro-scale purified antibodies in a total volume of 40 μL using DMEM supplemented with 2% FBS as a diluent and incubated for 1 h at 37°C in 5% CO2. At ∼17–20 h after seeding the cells, the virus–mAb mixtures were added to the cells in 384-well E-plates. Wells containing virus only (in the absence of mAb) and wells containing only Vero cells in medium were included as controls. Plates were measured every 8– 12 h for 48–72 h to assess virus neutralization. Micro-scale antibodies were assessed in four 5- fold dilutions (starting from a 1:20 sample dilution), and their concentrations were not normalized. In some experiments, mAbs were tested in triplicate using a single (1:20) dilution. Neutralization was calculated as the percent of maximal cell index in control wells without virus minus cell index in control (virus-only) wells that exhibited maximal CPE at 40 to 48 h after applying virus–antibody mixture to the cells. A mAb was classified as fully neutralizing if it completely inhibited SARS-CoV-2-induced CPE at the highest tested concentration, while a mAb was classified as partially neutralizing if it delayed but did not fully prevent CPE at the highest tested concentration.

#### Conventional throughput neutralization assay

To determine neutralizing activity of serum/plasma and IgG, we used real-time cell analysis (RTCA) assay on an xCELLigence RTCA MP Analyzer (ACEA Biosciences Inc.) that measures virus-induced cytopathic effect (CPE) (Gilchuk et al., 2020; Zost et al., 2020b). Briefly, 50 μL of cell culture medium (DMEM supplemented with 2% FBS) was added to each well of a 96-well E-plate using a ViaFlo384 liquid handler (Integra Biosciences) to obtain background reading. A suspension of 18,000 Vero-E6 cells in 50 μL of cell culture medium was seeded in each well, and the plate was placed on the analyzer. Measurements were taken automatically every 15 min, and the sensograms were visualized using RTCA software version 2.1.0 (ACEA Biosciences Inc). VSV-S (0.01 MOI, ∼120 PFU per well) was mixed 1:1 with a dilution of serum/plasma or mAb in a total volume of 100 μL using DMEM supplemented with 2% FBS as a diluent and incubated for 1 h at 37°C in 5% CO2. At 16 h after seeding the cells, the virus-mAb mixtures were added in replicates to the cells in 96-well E-plates. For the biliverdin assay, biliverdin was added to the virus at a final concentration of 25 μM before addition to the antibody; similarly, polysorbate-80 was added to the virus at 0.02% before addition to the antibody. Triplicate wells containing virus only (maximal CPE in the absence of mAb) and wells containing only Vero cells in medium (no-CPE wells) were included as controls. Plates were measured continuously (every 15 min) for 48 h to assess virus neutralization. Normalized cellular index (CI) values at the endpoint (48 h after incubation with the virus) were determined using the RTCA software version 2.1.0 (ACEA Biosciences Inc.). Results are expressed as percent neutralization in a presence of respective mAb relative to control wells with no CPE minus CI values from control wells with maximum CPE. RTCA IC50 values were determined by nonlinear regression analysis using Prism software.

Electron microscopy sample and grid preparation, imaging and processing of S6Pecto–Fab complexes. For electron microscopy imaging of spike protein and Fabs, we expressed a variant of S6Pecto containing a C-terminal Twin-Strep-tag, similar to that described previously (Zost et al., 2020b). Expressed protein was incubate with BioLock (IBA Lifesciences) and then isolated by Strep affinity chromatography on StrepTrap HP columns (GE Healthcare). Fabs were expressed as a recombinant Fab and purify with affinity column. For screening and imaging of negatively-stained SARS-CoV-2 S6Pecto protein in complex with human Fabs, the proteins were incubated at a Fab:spike molar ratio of 4:1 for about 1 hour at ambient temperature or overnight at 4°C, and approximately 3 μL of the sample at concentrations of about 10 to 15 μg/mL was applied to a glow-discharged grid with continuous carbon film on 400 square mesh copper electron microscopy grids (Electron Microscopy Sciences). The grids were stained with 0.75% uranyl formate (Ohi et al., 2004). Images were recorded on a Gatan US4000 4k×4k CCD camera using an FEI TF20 (TFS) transmission electron microscope operated at 200 keV and control with Serial EM (Mastronarde, 2005). All images were taken at 50,000× magnification with a pixel size of 2.18 Å per pixel in low-dose mode at a defocus of 1.5–1.8 μm. The total dose for the micrographs was around 30e−per Å2. Image processing was performed using the cryoSPARC (Punjani et al., 2017) software package. Images were imported, CTF-estimated and particles were picked. The particles were extracted with a box size of 256 pixels and binned to 128 pixels (pixel size of 4.36 A/pix) and 2D class averages were performed (see also Supplementary Table 3 for detailed). For time point of the complex with Fab Cov2-3434, SARS-CoV-2 S6Pecto protein and the Fab was mixed at ambient temperature and samples of ∼3 μL were pulled at the time points and applied to the grid and stained.

#### Serum antibody competition binding ELISAs with biotinylated reference mAbs

mAb COV2-3434 was biotinylated using NHS-PEG4-biotin (Thermo Fisher Scientific, cat# A39259) according to manufacturer protocol. Following biotinylation, biotinylated COV2-3434 was titrated in ELISA to verify specific binding and verify if EC50 was similar to the un-biotinylated antibody. Serum samples for use in competition ELISA were heat inactivated by incubation at 55°C for 1 hr. ELISAs were performed using 384-well plates that were coated overnight at 1 µg/mL with S6Pecto containing a C-terminal Twin-Strep-tag, similar to that described previously (Zost et al., 2020b). The following day, plates were washed three times with PBS-T and blocked with 2% bovine serum albumin (BSA) in PBS containing 0.05% Tween-20 (blocking buffer). Plates were washed three times with PBS-T and two-fold serial dilutions of donor serum (1:10 initial dilution) or control mAb (20,000 ng/mL initial dilution) in blocking buffer were added to each plate (total volume 25 µL/well) and incubated at RT for 1 hr. After incubation, 5 µL of biotinylated COV2-3434 (20 µg/mL) in blocking buffer were added directly to the wells containing the serial dilutions of competing serum or COV2-3434 mAb. The concentration of biotinylated mAb was calculated to be at approximately the EC90 of the mAb after addition to an equal volume of competing serum or mAb in the plate. Plates were incubated for 30 min at RT and then washed three times with PBS-T. After this wash, HRP-conjugated avidin (Sigma Aldrich, 1:3,500 dilution) in blocking buffer was added and plates were incubated for 1 h. After incubation, plates were washed three times with PBS-T and 25 µL of a 3,3′,5,5′ - tetramethylbenzidine (TMB) substrate (Thermo Fisher Scientific) was added to each well. After sufficient development, the reaction was quenched by addition of 25 µL 1 M HCl and the optical density values were measured at 450 nm wavelength on a BioTek plate reader. For each plate, background signal (signal from wells that were not coated with antigen) was subtracted and values were normalized to no-competition controls (signal from wells that had no competing serum or mAb) Four-parameter dose-response/inhibition curves were fit to the normalized data using Prism software (GraphPad) v8.1.1. Each dilution of serum or mAb was performed in triplicate and each experiment was conducted at least twice independently.

#### Protection against SARS-CoV-2 in mice

Animal studies were carried out in accordance with the recommendations in the Guide for the Care and Use of Laboratory Animals of the National Institutes of Health. The protocols were approved by the Institutional Animal Care and Use Committee at the Washington University School of Medicine (Assurance number A3381-01). Virus inoculations were performed under anesthesia that was induced and maintained with ketamine hydrochloride and xylazine, and all efforts were made to minimize animal suffering.

Female heterozygous K18-hACE C57BL/6J mice were housed in groups of up to 5 mice per cage at 18 to 24°C ambient temperatures and 40 to 60% humidity. Mice were fed a 20% protein diet (PicoLab 5053, Purina) and maintained on a 12-h light–dark cycle (06:00 to 18:00). Food and water were available ad libitum. Mice (8 to 9 weeks old) were inoculated with 1 × 10^4^ focus forming units of SARS-CoV-2 (viral titer was determined on Vero-TMPRSS2-ACE2 cells) via the intranasal route. Anti-SARS-CoV-2 human mAbs or isotype control mAbs were administered 24 h before (prophylaxis) SARS-CoV-2 inoculation. Weights and lethality were monitored daily for up to 6 days after inoculation and mice were euthanized at 6 dpi and tissues were collected.

#### Epitope mapping of antibodies by alanine-scanning mutagenesis

Epitope mapping was performed essentially as described previously (Davidson and Doranz, 2014) using a SARS-CoV- 2 (strain Wuhan-Hu-1) spike protein NTD shotgun mutagenesis mutation library, made using a full-length expression construct for spike protein, where 215 residues of the NTD (between spike residues 9 and 310) were mutated individually to alanine, and alanine residues to serine. Mutations were confirmed by DNA sequencing, and clones arrayed in a 384-well plate, one mutant per well. Binding of mAbs to each mutant clone in the alanine scanning library was determined, in duplicate, by high-throughput flow cytometry. A plasmid encoding cDNA for each spike protein mutant was transfected into HEK-293T cells and allowed to express for 22 h. Cells were fixed in 4% (v/v) paraformaldehyde (Electron Microscopy Sciences), and permeabilized with 0.1% (w/v) saponin (Sigma-Aldrich) in PBS plus calcium and magnesium (PBS++) before incubation with mAbs diluted in PBS++, 10% normal goat serum (Sigma), and 0.1% saponin. MAb screening concentrations were determined using an independent immunofluorescence titration curve against cells expressing wild-type S protein to ensure that signals were within the linear range of detection. Antibodies were detected using 3.75 μg/mL of Alexa-Fluor-488-conjugated secondary antibodies (Jackson ImmunoResearch Laboratories) in 10% normal goat serum with 0.1% saponin. Cells were washed three times with PBS++/0.1% saponin followed by two washes in PBS, and mean cellular fluorescence was detected using a high-throughput Intellicyte iQue flow cytometer (Sartorius). Antibody reactivity against each mutant S protein clone was calculated relative to wild-type S protein reactivity by subtracting the signal from mock-transfected controls and normalizing to the signal from wild-type S-transfected controls. Mutations within clones were identified as critical to the mAb epitope if they did not support reactivity of the test MAb but supported reactivity of other SARS-CoV-2 antibodies. This counter-screen strategy facilitates the exclusion of S protein mutants that are locally misfolded or have an expression defect.

#### Measurement of viral burden

Plaque assays were performed as described previously (Case et al., 2020; Hassan et al., 2020) on Vero+TMPRSS2+hACE2 cells. Briefly, lung homogenates were serially diluted and added to Vero+TMPRSS2+hACE2 cell monolayers in 12-well plates. Plates were incubated at 37°C for 1 h and then overlaid with 1% (w/v) methylcellulose in MEM supplemented with 2% FBS. Plates were incubated at 37°C for 1 h and then overlaid with 1% (w/v) methylcellulose in with 4% PFA for 20 min. Plaques were visualized by staining with 0.05% crystal violet in 20% methanol.

#### Quantification and statistical analysis

Mean ± S.E.M. or mean ± S.D. were determined for continuous variables as noted. Technical and biological replicates are described in the figure legends. For analysis of mouse studies, the comparison of weight-change curves was performed using a one-way ANOVA with Dunnett’s post hoc test of the area under the curve for days 3-6 post-infection, using Prism v.9.0 (GraphPad). Infectious viral loads were compared by a one-way ANOVA with Dunnett’s multiple comparisons test using Prism v.9.0 (GraphPad).

**Figure S1. Divergence from inferred germline gene sequences, related to Figure 2**

A. The number of mutations of each mAb relative to the inferred germline variable gene was counted for each clone. These numbers then were transformed into percent values and plotted as violin plots. For the heavy chain, values range from 80.4 to 100, with a median of 96.6, a 25th quartile of 94.6 and a 75th quartile of 97.6. For the light chain, values range from 88.1 to 100, with a median of 97.9, a 25th quartile of 96.5 and a 75th quartile of 98.9.

B. Bar graph showing IGHV gene usage by 102 clones expressed Y-axis represents number of times same IGHV appeared and on x-axis is the IGHV gene identified.

**Figure S2. Gating strategy for cell-surface antigen-display experiment, related to Figure 3**

A. The first gate is for all cells, the second gate is for infected cells, and the third gate is for antibody binding to infected cells.

B. Overlay of histograms infected cells in light grey on uninfected cells in dark grey gated for Alexa Fluor 647 staining.

**Figure S3. Phylogenetic tree obtained after aligning multiple sequence of the heavy chain of IGHV1-24 genes, related to Figure 3** identified in this study (red) with IGHV 1-24 genes available from sequences 1) deposited in public databases shown in cyan color, 2) from vaccinated individuals shown in purple, or 3) from infected individuals shown in orange.

**Figure S4. Competition ELISA of mAbs, related to Figure 4**

Competition ELISA of mAbs with previously mapped antibodies COV2-2130, COV2-2196, COV2-2676, COV2-2489, r4A8 or rCR3022. Unlabeled antibodies applied to antigen first are indicated on the left, while biotinylated antibodies that were added to antigen-coated wells second are listed across the top. The number in each box represents the percent competition binding of the biotinylated antibody in the presence of the indicated competing antibody. Heat map colors range from dark grey (100% blocking of the biotinylated antibody) to white (0% or no blocking of the biotinylated antibody).

**Figure S5. Epitope identification and characterization of COV2-3439, related to Figure 4**

Residues critical for COV2-3439 binding, identified by screening COV2-3439 on an NTD alanine-scan mutagenesis library, are shown in red spheres on the NTD (PDB 7L2C).

**Figure S6. Structural characterization of COV2-3434, related to Figure 5**

Steric clash of COV2-3434 Fab (green) with SARS-CoV2- S monomer (cyan) in open conformation when modeled double Fab (COV2-3434 Fab (green) COV2-3439 Fab (magenta) + rNTD (blue) complex on to SARS-CoV2- S monomer (cyan) in open conformation.

**Figure S7. Neutralization of VSV-S by COV2-3434 related to Figure 5**

A. Neutralization of VSV-S by COV2-3434 was measured in the absence or presence of 0.02% polysorbate-80 in Vero-CCL81 cells.

B. Neutralization of VSV-S by COV2-3434 was measured in the absence or presence of 25 μM biliverdin in Vero-CCL81 cells.

**Figure S8. Protection in K18 hACE2 transgenic mice by trimer-disrupting antibody COV2- 3434, related to Figure 6.**

Eight-week-old female K18-hACE2 transgenic mice were inoculated by the intranasal route with 104 FFU of SARS-CoV-2 (WA1/2020 D614G). One day prior to virus inoculation, mice were given a single 1 mg dose of COV2-3434, COV2-2196, or isotype control mAb by intraperitoneal injection. Data are from two independent experiments, n=7 (isotype) or 8 (all other groups).

A. Weight was monitored daily. Two-way ANOVA with Dunnett’s post-test with comparison to control mAb: **, p<0.001; *, p<0.05; ns, not significant.

B. At 6 dpi, tissues were collected, and viral RNA levels in indicated tissues were determined (line indicates median). One-way ANOVA with Dunnett’s post-test: ****, p<0.0001; *p<0.05; ns, not significant. The dotted line represents the limit of detection (LOD) of the assay.

