## Supplemental Information for "An antibody targeting the N-terminal domain of SARS-CoV-2 disrupts the spike trimer"

**A**

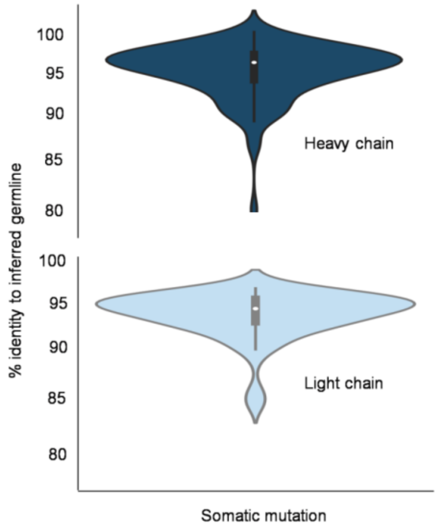


**B**

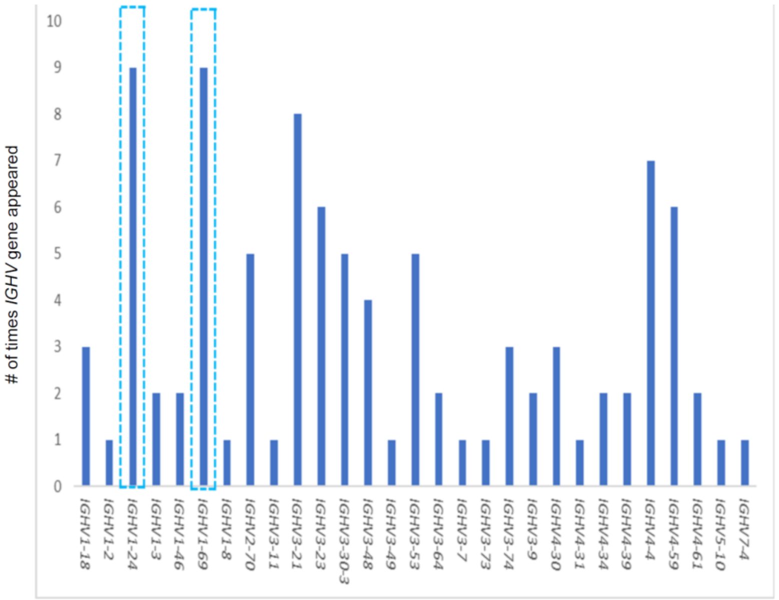


**Figure S1. Divergence from inferred germline gene sequences, related to Figure 2**

A

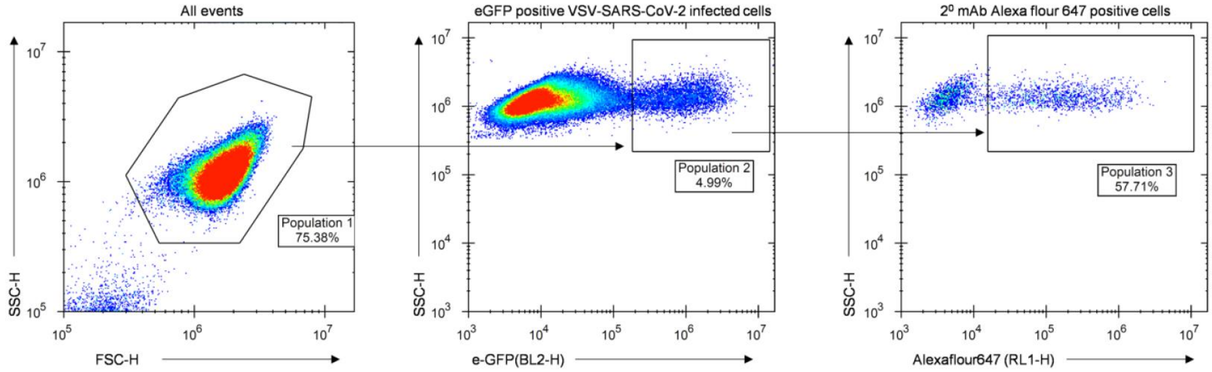


B

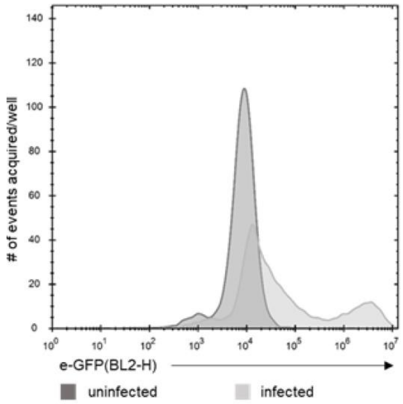


**Figure S2. Gating strategy used for cell-surface antigen-display experiment, related to Figure 3**

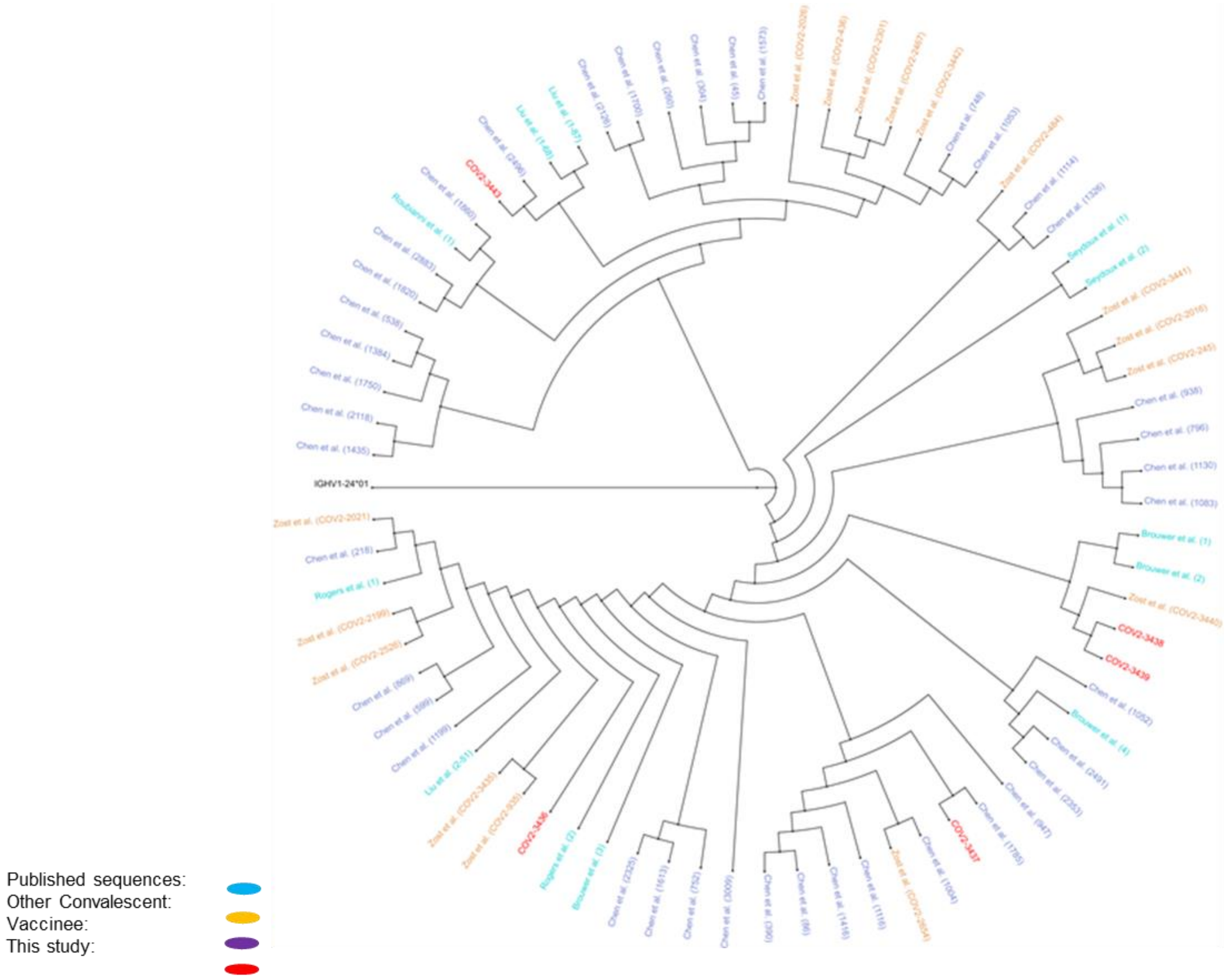


**Figure S3. Phylogenetic tree obtained after aligning multiple sequence of the heavy chain of *IGHV1-24* genes, related to Figure 3**

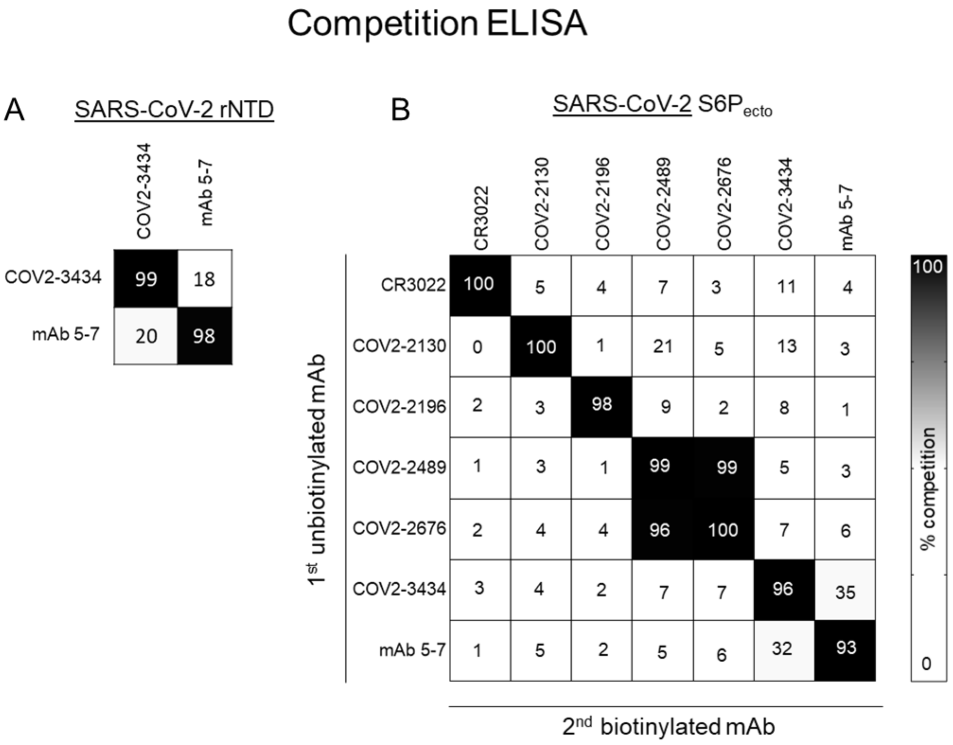


**Figure S4. Competition ELISA of mAbs, related to Figure 4**

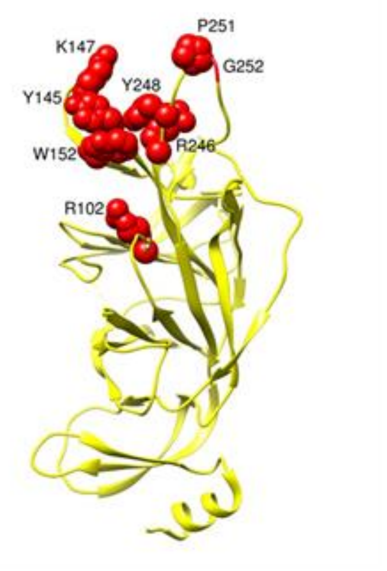


**Figure S5. Epitope identification and structural characterization of COV2-3439, related to Figure 4**

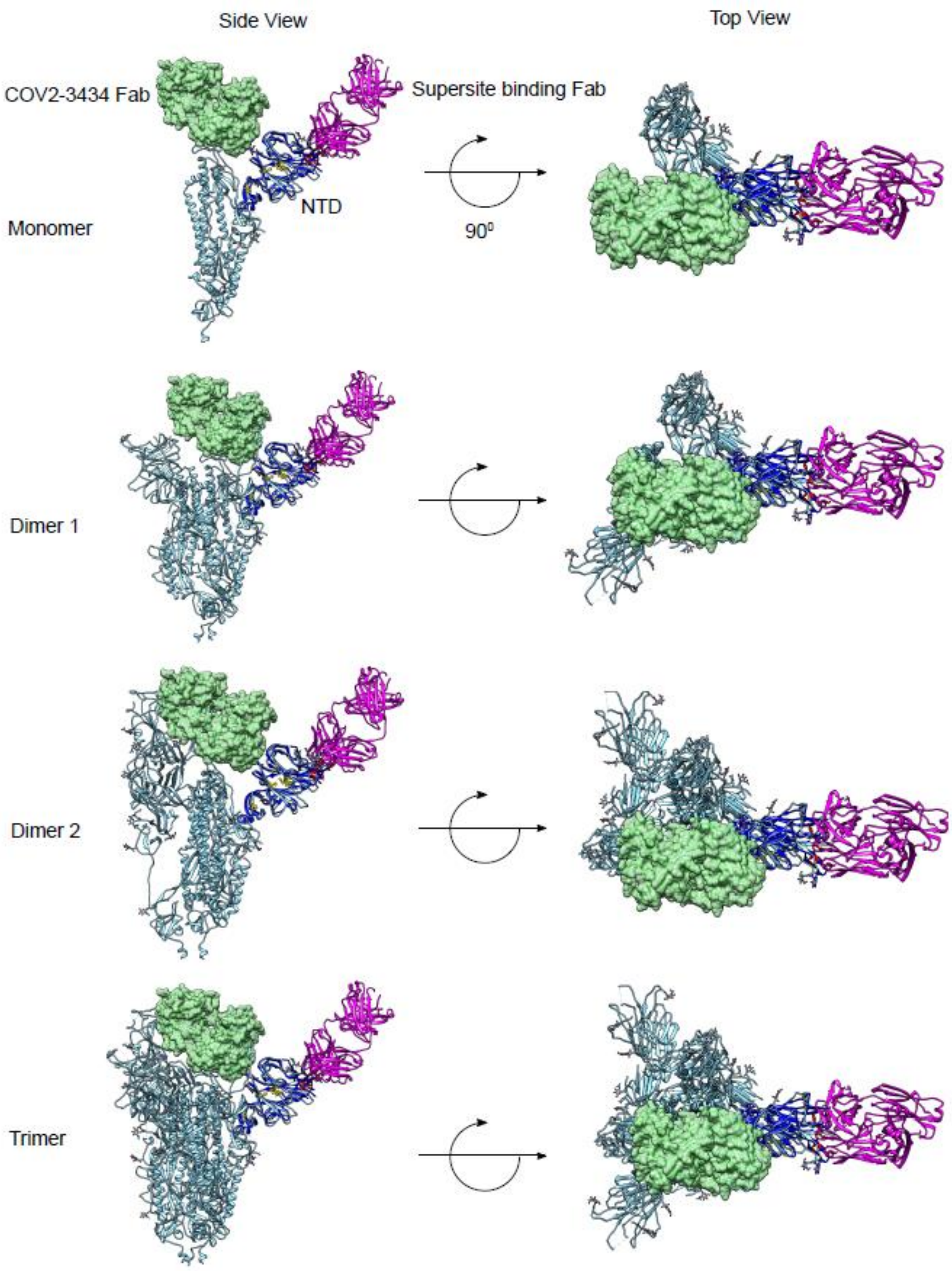


**Figure S6. Structural characterization of COV2-3434, related to Figure 5**

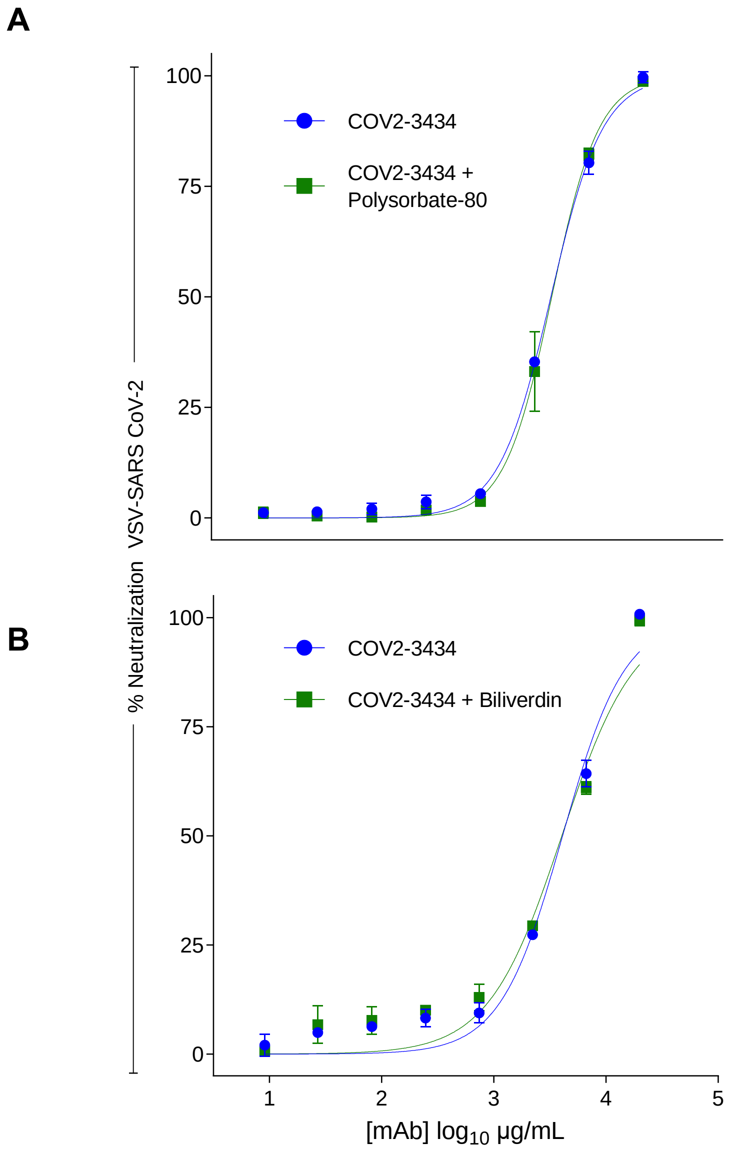


**Figure S7. Neutralization of VSV-S by COV2-3434 related to Figure 5**

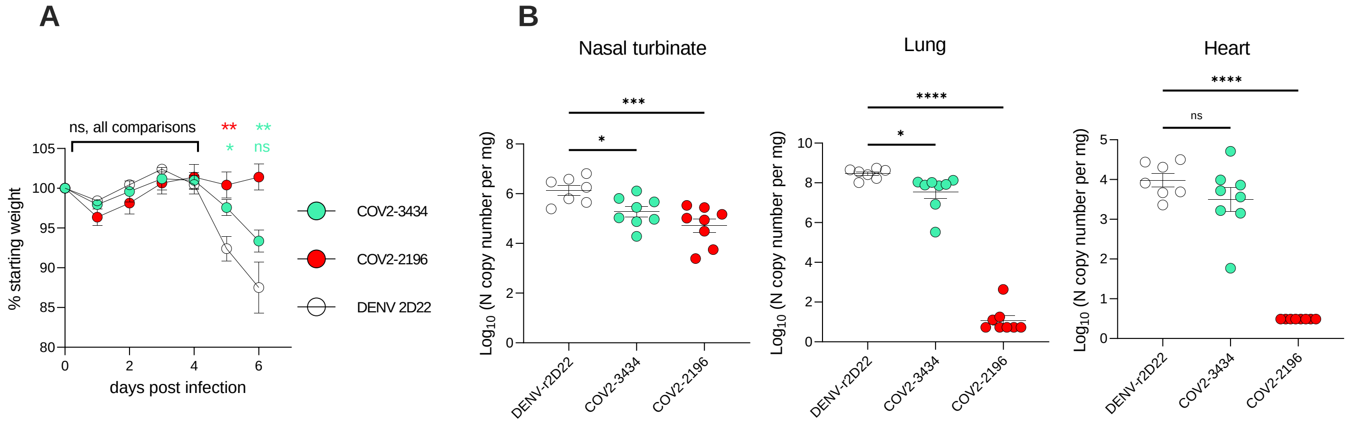


**Figure S8. Protection in K18 hACE2 transgenic mice by trimer-disrupting antibody COV2-3434, related to Figure 6.**

**Table S1. Clinical features of individuals studied**

| **Exposure** | **Donor #** [vaccine (# doses)] | **Location of exposure** | **Sample from timepoint after exposure (days)** | **Symptoms** |
|---|---|---|---|---|
| **Infection** | 1989 | Nashville, TN, USA | 28 | Fever, chills, fatigue, cough, shortness of breath, loss of appetite |
| | | | 56 | |
| | | | 112 | |
| | 1988 | Nashville, TN, USA | 16 | Fever, fatigue, cough, shortness of breath, loss of appetite |
| | | | 28 | |
| | | | 56 | |
| | 1992 | Nashville, TN, USA | 13 | Fever, chills, fatigue, cough, shortness of breath, headache, myalgia |
| | | | 28 | |
| | | | 56 | |
| | 1951 | Beijing, China | 35 | Low-grade fever, cough, runny nose |
| | | | 36 | |
| | | | 75 | |
| **Vaccination** | 269 [Pfizer (2)] | Nashville, TN, USA | *pre-vac* -24 | NA |
| | | | *post-vac* 250 | NA |
| | 1672 [Pfizer (2)] | Nashville, TN, USA | *pre-vac* -327 | NA |
| | | | *post-vac* 233 | NA |
| | 1801 [Pfizer (2)] | Nashville, TN, USA | *pre-vac* -773 | NA |
| | | | *post-vac* 10 | NA |
| **Vaccination followed by breakthrough infection** | 1079 [Pfizer (1)] | Nashville, TN, USA | *pre-vac* -111 | NA |
| | | | *post-vac* 193<br>*post-inf* 66 | Fever, chills, fatigue, cough, shortness of breath, headache, myalgia, loss of smell, loss of taste |
| | 2062 [Pfizer (2)] | Nashville, TN, USA | *pre-vac* -11 | NA |
| | | | *post-vac* 201<br>*post-inf* 34 | Fever, cough, loss of smell, headache, sore throat |

NA, indicates Not applicable.

**Table S2**. *IGHV1-24* genes identified in our study or from published data, with CDRH3 amino acid length and sequences.

| Publication | Monoclonal antibody | *IGHV* | CDRH3 length (# amino acids) | CDRH3 amino acid sequence | *IGLV* |
|---|---|---|---|---|---|
| Chi X *et al.*, 2020 | 4A8 | *IGHV1-24* | 21 | ATSTAVAGTPDLFDYYYGMDV | *IGKV2-24* |
| Liu L *et al.*, 2020 | 1-68 | ” | 21 | ATGWAVAGSSDVWYYYYGMDV | *IGLV2-18* |
| Liu L *et al.*, 2020 | 1-87 | ” | 21 | ATGIAVIGPPPSTYYYYGMDV | *IGLV2-14* |
| This study | COV2-3443 | ” | 21 | ATAIAVAGSPEYYYYYHGMDV | *IGLV1-44* |
| This study | COV2-3438 | ” | 14 | AISPAIVAAGWLDP | *IGLV2-23* |
| Voss *et al.*, 2021 | CM25 | ” | 14 | ATGPAVRRGSWFDP | *IGLV1-51* |
| Liu L *et al.*, 2020 | 2-51 | ” | 14 | ATGWAYKSTWYFGY | *IGLV2-8* |
| Zhang L *et al.*, 2020 | FC05 | ” | 14 | ATTTPFSSSYWFDP | *IGLV1-51* |
| This study | COV2-3436 | ” | 14 | ATVFAIFGVVRFDY | *IGLV1-40* |
| This study | COV2-3439 | ” | 15 | ATSSPIVGTTGWFDP | *IGKV1-39* |
| This study | COV2-3437 | ” | 12 | ATGHQLLVHWFDP | *IGKV3-20* |

| **Table S3. Summary of electron microscopy data collection and statistics SARS-CoV-2$_{ecto}$ protein in complex with human Fabs** | | | | |
|---|---|---|---|---|
| | | **Structure of SARS-CoV-2 S6P$_{ecto}$ protein in complex with indicated Fab** | | **Structure of SARS-CoV-2 rNTD in complex with** |
| | | **Fab COV2-3434** | **Fab COV2-3439** | **Fab COV2-3434/COV-3439** |
| **Microscope setting** | Microscope | TF-20 | TF-20 | TF-20 |
| | Voltage (kV) | 200 | 200 | 200 |
| | Detector | US-4000 CCD | US-4000 CCD | US-4000 CCD |
| | Magnification | 50,000x | 50,000x | 50,000x |
| | Pixel size | 2.18 | 2.18 | 2.18 |
| | Exposure (e-/Å2) | 30 | 30 | 30 |
| | Defocus range (µm) | 1.5 to 1.8 | 1.5 to 1.8 | 1.5 to 1.8 |
| **Data** | Micrographs, # | 319 | 320 | 300 |
| | Particles, # | 70,634 | 48,000 | 210,000 |
| | Particles #, after 2D | 16,012 | 33,360 | 90,476 |
| | Final particles, # | 16,012 | 18,375 | 90,476 |
| | Symmetry | C1 | C1 | C1 |
